# Restoring data balance via generative models of T cell receptors for antigen-binding prediction

**DOI:** 10.1101/2024.07.10.602897

**Authors:** Emanuele Loffredo, Mauro Pastore, Simona Cocco, Rémi Monasson

**Author notes:** **For correspondence:** (MP). These authors contributed equally to this work (Emanuele Loffredo, Mauro Pastore). These authors also contributed equally to this work (Simona Cocco, Rémi Monasson).

## Abstract

Unveiling specificity in T cell recognition of antigens represents a major step to understand the immune system response. Many supervised machine learning approaches have been designed to build sequence-based predictive models of such specificity using binding and non-binding receptor-antigen data. Due to the scarcity of known specific T cell receptors for each antigen compared to the abundance of non-specific ones, available datasets are heavily imbalanced and make the goal of achieving solid predictive performances very challenging. Here, we propose to restore data balance through data augmentation using generative unsupervised models. We then use these augmented data to train supervised models for prediction of peptide-specific T cell receptors, or binding pairs of peptide and T cell receptor sequences. We show that our pipeline yields increased performance in prediction tasks of T cell receptors specificity. More broadly, our pipeline provides a general framework that could be used to restore balance in other computational problems involving biological sequence data.

## Introduction

Performances of Machine Learning (ML) approaches are well known to crucially depend on the composition and quality of training data. In the case of classification, imbalance in the training dataset – which occurs when one or more classes are significantly under-represented – poses serious challenges during the learning process, as most algorithms are designed to optimize performances over all data, leading to biased models that mostly focus on the majority classes. Yet, the correct characterization of rare examples is of primarily relevance, in particular in many biologically-related applications, such as molecular biology (***Wang et al., 2006; Yang et al., 2012; Cheng et al., 2015; Song et al., 2021; Ansari and White***, ***2024***), automated medical diagnostics (***Krawczyk et al., 2016; Fotouhi et al., 2019***), ecology (***Kyathanahally et al., 2021; Chen et al., 2025***), etc. Among these, adaptive immune receptors data, which are the focus of this work, represent a typical scenario where machine learning approaches might be limited due to the imbalance: we will address this question in the context of predicting the specificity of T cell receptors (***Deng et al., 2023; Mason and Reddy, 2024***).

Cytotoxic T lymphocytes (T cells) play a crucial role in the adaptive immune response of organisms against pathogens and/or malfunctioning cells (***Zhang and Bevan, 2011***). Short pathogen protein regions (peptide antigens) interact with the Major Histocompatibility Complex proteins (MHC) and form peptide-MHC epitopes (pMHC). Binding of CD8+ T cell receptor (TCR) with pMHC enables the killer machinery against such pathogen. Given the high specificity of the interaction, TCRs bind a limited number of presented pMHC. Achieving a reliable prediction of TCR-pMHC binding from sequence data represents a major goal in the field, in particular for the development of vaccines and the improvement of personalized immunotherapies.

Over the recent years, much progress in this direction has been made with computational approaches (***Sim, 2024; Meysman et al., 2023; Ghoreyshi and George, 2023; Weber et al., 2023; Nagano et al., 2025***), benefiting both from the power of ML methods and the large-scale amount of experimentally tested data. Such works typically use the sequences of the Complementarity-Determining Region-3 beta (CDR3*β*) and alpha (CDR3*α*) chains paired with peptide sequences to reveal TCR-pMHC binding affinity. The CDR3 region is the most variable one in TCRs and is recognized to be the major actor influencing TCR specificity for peptide binding. Though recent works have shown that use of both *α* and *β* chains leads to better predictions (***Dash et al., 2017; Montemurro et al., 2021***), many works still focus on *β* chains solely, because they primarily drive the immune response (***Springer et al., 2021***) and are more abundant in most databases.

Predicting TCRs specificity is a computationally challenging problem for several reasons. On the one hand, CDR3*β* sequences binding different target epitopes exhibit very strong similarities and it is hard to identify the features determining binding specificity. On the other hand, ML predictive models are trained on labelled (experimentally tested) TCR-pMHC sequence data through supervised learning to obtain accurate predictions. Public databases containing TCR specificity, such as IEDB (***Vita et al., 2018***), VDJdb (***Shugay et al., 2017; Bagaev et al., 2019***), Mc-PAS TCR (***Tickotsky et al., 2017***) and PIRD (***Zhang et al., 2019***), mostly include limited amount of TCR sequences with positive interaction, *i*.*e*. known to bind some peptides of interest. Considerable efforts have been undertaken to build *ad hoc* negative sequence datasets, for instance by mismatch pairing and/or taking bulk (unlabelled) data from healthy donors. While the way the negative data are chosen may potentially bias predictions (***Ursu et al., 2024***), their number exceed in quantity, by several orders of magnitude, the amount of experimentally tested positive binding pairs. As a result, binding prediction models are trained from datasets with strong imbalance between the positive and negative classes.

The present work aims first at analyzing how much TCR specificity predictions are hindered by the imbalance issue. We then propose a two-step pipeline to restore data balance using a generative framework for specific TCR sequences. Our findings highlight that it is worth to carefully restore balance in the composition of training datasets in order to improve performances. Furthermore, since our pipeline is quite general and can handle various sequence-based data, its relevance reaches beyond the TCR field, to any sequence-based classification problem where imbalance is of concern (***Wang et al., 2006; Yang et al., 2012; Cheng et al., 2015; Song et al., 2021; Ansari and White, 2024***).

The paper is organized as follows. In Background and relevance, we present a general overview on the problem of learning with class imbalance. In Results, we introduce our pipeline (detailed in Materials and methods) to tackle this problem in the context of CDR3*β* binding predictions, and we give evidence on how it improves performances on both peptide-specific and pan-specific tasks. In Discussion and conclusions, we draw our conclusions.

## Background and relevance

### Effect of class imbalance on decision boundary

Class imbalance is a widespread problem affecting both bio- and non bio-related sources of data and is notably an Achilles’ heel in applying machine learning methods to achieve reliable predictions (***Fernández et al., 2018***). Informally speaking, naive ML models adapt to the most represented class in the data and are less accurate in capturing the features of under represented examples, degrading the quality of predictions whenever the minority class is of crucial interest. From a geometrical perspective, supervised architectures solve classification tasks by learning a decision boundary in the feature space of input data: the effect of class imbalance can be envisioned as shifting away the decision boundary from the optimal one, at the expenses of classes with fewer data points.

To visualize this effect in a simple setting, let us first focus on generalized linear architectures, which classify input points by learning separating hyperplanes. The high-dimensional (in data dimension and size of the training set) asymptotics of learning and generalization can be fully tackled theoretically within the framework of equilibrium statistical physics (***Del Giudice, P. et al., 1989; Loureiro et al., 2021b; Pesce et al., 2023; Dandi et al., 2023; Mannelli et al., 2023; Loffredo et al., 2024; Pezzicoli et al., 2025***). This analysis, which has been proven relevant in realistic cases (***Loureiro et al., 2021a; Pesce et al., 2023; Loffredo et al., 2024***), shows that classifiers trained on imbalanced datasets learn a sub-optimal decision hyperplane, with a systematic offset from the best attainable one, as well as errors on the orientation (Fig. 1a).

**Figure 1.**
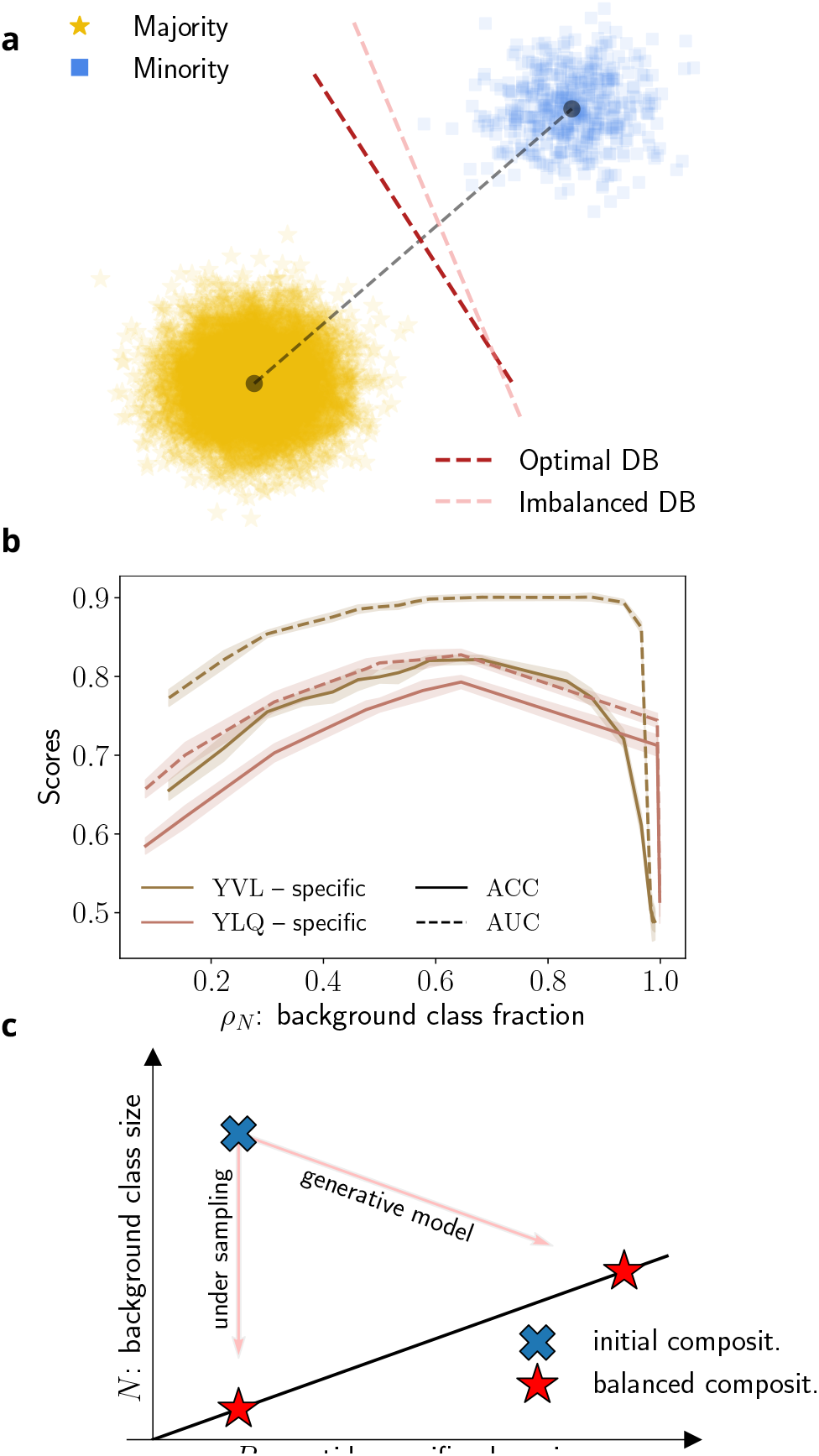
Effects of imbalanced data on ML classifiers. **a)** 2D visualization of a two-class training dataset, with the majority class (yellow) significantly outnumbering the minority one (blue). Data are generated from a Gaussian mixture; the decision boundary (DB) separating the classes passes perpendicularly through the midpoint of the line segment connecting the centers of the two clusters (dark line); a linear classifier trained under imbalance learns a sub-optimal decision boundary (light line), which leads to low predictive capabilities (see App. 2 for more details). **b)** Accuracy and AUC scores for a predictive model trained to distinguish `YVLDHLIVV`- and `YLQPRTFLL`-specific CDR3*β* sequences from bulk CDR3*β*s as a function of the fraction *ρ*_*N*_ of background data in the training set. In practice we fix the class size of peptide-specific sequences and vary the size of the background sequences class to change *ρ*_*N*_. Performances (evaluated on a balanced test set) are optimal when the two class sizes are of roughly equal sizes, *i*.*e*. when *ρ*_*N*_ ≃ 0.5. **c)** Graphical visualization of the imbalanced composition of TCRs datasets, in a two-class setting where *P, N* represent the class sizes. Our work proposes to restore class balance (*i*.*e. P* = *N*, straight line) by introducing a generative model able to sample new sequences compatible with the positive class, for which few data are experimentally available.

This picture qualitatively holds for deeper architectures, for two essential reasons: (i) the last layer of a deep classifier is usually fully-connected and can be thought of as a linear classifier in the space of the features learned by the previous layers; (ii) in certain regimes, such as the infinite-width limit and lazy training (***Neal, 1996; Williams, 1996; Lee et al., 2018; de G. Matthews et al., 2018; Jacot et al., 2018; Bietti and Mairal, 2019; Chizat et al., 2019***), deep models have been proven equivalent to kernel machines, which embed the data in a randomized feature space and operate there as linear classifiers (***Dietrich et al., 1999; Gerace et al., 2021; Aguirre-López et al., 2025***).

### Metrics for performance assessment

An important issue is how to fairly assess performance under imbalance. Consider for instance a dataset with two classes having imbalance ratio 1:9 in the data. Suppose we split the data along a canonical training/test cross-validation procedure, maintaining on average the same imbalance ratio in each subset, and that we naively assess performances using the standard measure of accuracy – which counts the number of correctly classified examples in the test set. Even a null model identifying everything as the majority class would have high accuracy (≈ 90%), yet such result is clearly meaningless in quantifying how good the model can distinguish the two data classes. Similar warped assessments are obtained from other evaluation metrics that explicitly depend on the test set composition. Furthermore, imbalance can have an implicit and more subtle impact on evaluation metrics insensitive to some model parameters that are however used at inference level. This is the case of measures obtained by thresholding the predictor of the model and integrating parametric curves of the threshold, such as the area under the Receiver Operating Characteristic curve, or the area under the Precision-Recall curve. These estimators are not sensitive to the bias of the model (the offset of the hyperplane for linear classifiers in Fig. 1a), which is often the parameter most impacted by imbalance during training (***Loffredo et al., 2024***).

Many studies have discussed the most useful metrics to adopt in this context (***Forman and Scholz, 2010***), with no clear agreement in the literature on which one is less prone to imbalance biases (***Lobo et al., 2008; Saito and Rehmsmeier, 2015; Richardson et al., 2024***). Here, we follow the recent work ***Loffredo et al. (2024)***, where a theoretical comparison of different evaluation metrics was performed, suggesting that it is best to use metrics that do not depend explicitly on the imbalance on the test set, i.e. that equally weight the majority and minority contributions regardless of the imbalance ratio, or to use an explicitly balanced test set. Consequently, we hereafter evaluate model performances measuring the accuracy (ACC) and Area Under the Receiver Operating Characteristic curve (AUC) always on a balanced test set, regardless of the training set imbalance (see details in Materials and methods). In Fig. 1b we plot the behavior of these metrics for models trained to classify CDR3*β* binders to two different epitopes from background sequences with varied imbalance ratios. We observe that the best performance are, not surprisingly, obtained when the training data are (close to) being balanced.

### Mitigation approaches

A plethora of mitigation strategies have been proposed to cure the effect of imbalance in the training data and improve performances. Some of these approaches act at the algorithmic level (***Fernández et al., 2018)***, and proceed by reweighting the loss (to boost minority data), adapting the optimizer to account for imbalance, or averaging over ensembles (e.g. bagging), etc… Data-level methods seek to restore data balance through downsampling majority data or data augmentation of the minority class. Here, we focus on the latter set of methods for mainly two reasons: first, simple algorithmic strategies such as class reweighting in the training loss can be thought as equivalent to random oversampling of the minority class; second, developing generative models to augment data has an interest *per se* for the design of synthetic sequences. In practice, starting with a given dataset composition with two classes – the majority and minority one – we combine majority undersampling and minority oversampling in order to reach some intermediate common size for both classes, see Fig. 1c and Materials and methods for details. In a recent theoretical study, we have benchmarked different strategies to produce a balanced dataset (***Loffredo et al., 2024)***, and showed that such a mixed approach offered the highest predictive performances. In this work we will follow this line, showing that also for sequence-based problems and specifically for TCR specificity predictions, mixed strategies yield better results.

## Results

### Learning pipeline

We now present our mixed mitigation strategy, which combines data removal (subsampling of the negative class) and, crucially, data augmentation (oversampling of the positive class), see Fig. 1c. We implement this mixed strategy for both

1. *peptide-specific* models, trained on one or more selected epitopes and for which the peptide sequence is only used as a label during the training. This setting corresponds to multi-class classification of CDR3*β* sequences, with classes identified by the binding peptides.
2. *pan-specific* models, where combinations of peptides and TCR sequences are presented during training, together with binary labels expressing if the peptide-TCR pairs are binding or not. This setting corresponds to binary classification of CDR3*β* + peptide binding/non-binding pairs.

Though pan-specific models are harder to build as they require more data than their peptide-specific counterparts, they can, in principle, leverage the diversity of the peptide space to capture the common underlying features of TCR-peptide interactions and potentially recognize binding to new, unseen antigens. We explore this last possibility by studying the out-of-distribution performances of our predictive models.

The learning pipeline, fully presented in Fig. 2 for both peptide- and pan-specific cases, consists of two steps. First, data balance is restored by randomly subsampling the negative data class and by augmenting the positive class of sequences through data generation. To do so, we learn an unsupervised generative model over the limited number of peptide-specific CDR3*β* sequences in the training set and enlarge the positive class by generating surrogate sequences. The idea of using unsupervised generative model for data augmentation to mitigate imbalance or data scarcity was recently proposed in a variety of settings (***Zięba et al., 2015; Wan et al., 2017; Douzas and Bacao, 2018; Mirza et al., 2021; Mondal et al., 2023; Ai et al., 2023; Loffredo et al., 2024)***, and was successfully applied to biological-sequences related tasks, *e*.*g*. in ***Marouf et al. (2020)***. We consider two generative models, namely

**Figure 2.**
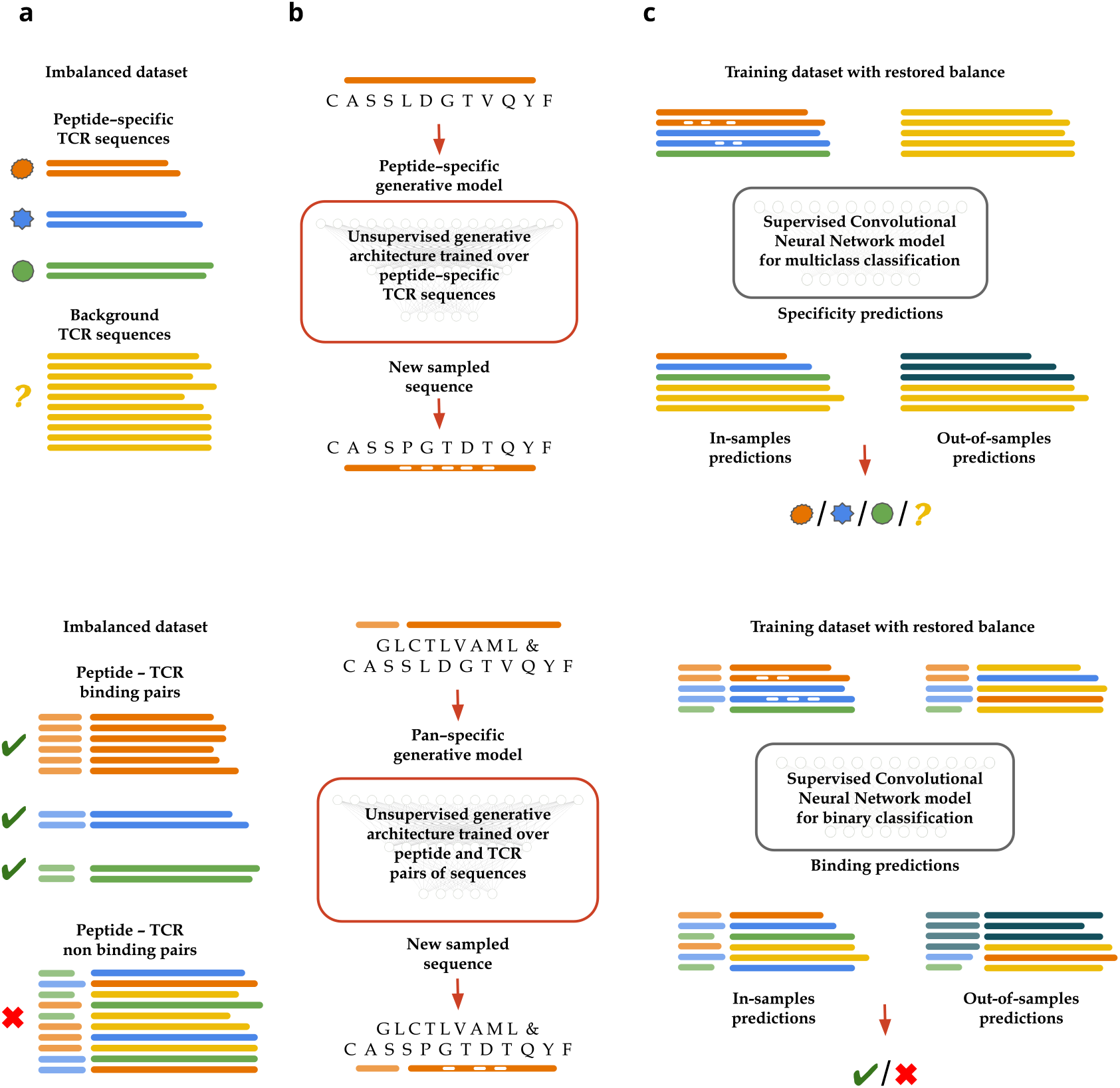
Learning pipelines for peptide-specific (top) and pan-specific (bottom) models. **(Top)**: peptide-specific models. Column **a)**: Data consists of few CDR3*β* sequences, known to bind some epitopes (colored symbols and segments) and of many ‘negative’ sequences (yellow). Column **b)**: A generative model is trained over peptide-specific CDR3*β* sequences, here, corresponding to the orange epitope. After training, Gibbs sampling of the inferred probability landscape allows us to generate putative peptide-specific sequences. Column **c)**: A supervised CNN architecture is trained over (natural and generated) peptide-specific CDR3*β*s and background CDR3*β*s; after learning, the network is used as predictive model for TCR specificity over in- and out-of-distribution (black sequences) test data. **(Bottom)**: Pan-specific models. Column **a)**: Compared to the pipeline above, input data are joint sequences of peptides (left, lighter color) and of TCR (right). Background sequences are obtained through mismatch pairing. Column **b)**: The generative models produce putative binding pairs of peptide and TCR sequences. Column **c)**: Supervised classifier trained to carry out TCR-epitope binding predictions.

- Restricted Boltzmann Machines (RBMs), two-layer architectures extracting latent features from sequence data;
- Bidirectional Encoder Representation Transformer (BERT)-like architectures trained over CDR3*β* sequences to learn their *grammatical structure*.

For peptide-specific predictions we separately train a model for each epitope, while for pan-specific predictions a collective model encompassing all the epitopes receives as input peptide and CDR3*β* data, see Generative models for data augmentation.

Second, we use the natural and generated data to train a supervised model for TCR-epitope binding predictions. Models existing so far range from random forests (***Gielis et al., 2019; Pham et al., 2023)*** to neural network architectures of different complexities, *e*.*g*. convolutional neural networks (***Montemurro et al., 2021; Sidhom et al., 2021)***, long-short term memory networks and autoencoders (***Springer et al., 2020; Lu et al., 2021)***; unsupervised algorithms have also been adapted to this task, such as SONIA (***Sethna et al., 2020)*** and its more precise variant soNNia (***Isacchini et al., 2021)***, diffRBM (***Bravi et al., 2023)***, which implements transfer learning within Restricted Boltzmann Machines, and, in the context of Large Language Models, Transformers-based approaches (***Yadav et al., 2024; Wu et al., 2024; Meynard-Piganeau et al., 2024)***.

Hereafter, we resort to a one-dimensional convolutional neural network (CNN) architecture, trained over positive (natural and generated) and negative sequences – once balance has been restored between classes. The architectures for peptide- and pan-specific models are slightly different, see Fig. 2, and are detailed in Model architecture for supervised predictions. Last of all, the predictive power of the model is tested over its ability to discriminate positive against negative CDR3*β* sequences (the in-distribution test set) or against other new peptide-specific CDR3*β* (the out-of-distribution test set).

### Peptide-specific models for in-distribution predictions benefit from data augmentation

We present below results for a case where the predictive model has to learn three different classes of peptide-specific receptors (labelled with *p*_1_, *p*_2_ and *p*_3_) and bulk receptors (referred to as *b*). To assess if an unsupervised model that generates CDR3*β* sequences to augment the peptide-specific classes size can yield better performances than naive undersampling of each CDR3*β* class down to the lowest available class size, we consider two different strategies to restore balance in the dataset:

- A first protocol, in which data are unaltered and each class size 𝒟 is given by

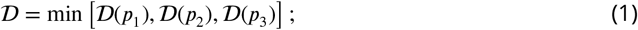

*i*.*e*. data points in over-represented classes are randomly under sampled down to the common size 𝒟.
- A second protocol, in which small-size classes are augmented using the generative model pipeline up to a target size 𝒢 common to all classes, with 𝒢 ≤ 10 𝒟 for computational reasons.

Notice that, for both protocols, the bulk class size 𝒟 (*b*), which is orders of magnitude larger than any 𝒟 (*p*_*i*_), is randomly under sampled to match the final common size of the positive data. We use as generative framework both a RBM- and a BERT-based sampling strategy, to assess the effects of balancing through data augmentation regardless of the generative model. These data are then used to train our classifier. Notice that the last layer of the CNN architecture, which outputs the binding predictions, is designed to have as many units as the number of classes, *i*.*e*. of peptide labels plus the background label, with softmax activation function. This allows us to carry out multiclass classification, as the network is able to predict specificity towards more than one target.

Fig. 3 shows results for multiple experiments related to different triplets of peptide-specific CDR3*β* sequences, confirming that the quality of predictions increases with the pipeline depicted in Fig. 2 (top row). In particular, the gain in performance is larger for multiclass experiments where one (or more) class contains few sequences, *e*.*g*. 𝒟 (*p*_1_) ≪ 𝒟 (*p*_2_), 𝒟 (*p*_3_). For example, experimental data for the epitopes `AMFWSVPTV`, `VTEHDTLLY` and `GLCTLVAML` have relative sizes 1.5% − 2.4% − 96.1%, which introduces a strong bias towards the `GLCTLVAML`-specific CDR3*β*s. Restoring balance by generating new `AMFWSVPTV`-specific CDR3*β* sequences, we are able to obtain a 20% performance increase compared to restoring balance by undersampling only. This gain is milder when pepitde-specific classes have more sequences and are more balanced among each other, as the case of epitopes `ELAGIGILTV` (9%), `LLWNGPMAV` (21%) and `GLCTLVAML` (70%) shows. Relative abundances of the initial dataset for the combinations reported in Fig. 3 can be obtained from the absolute class sizes reported in App. 1.

**Figure 3.**
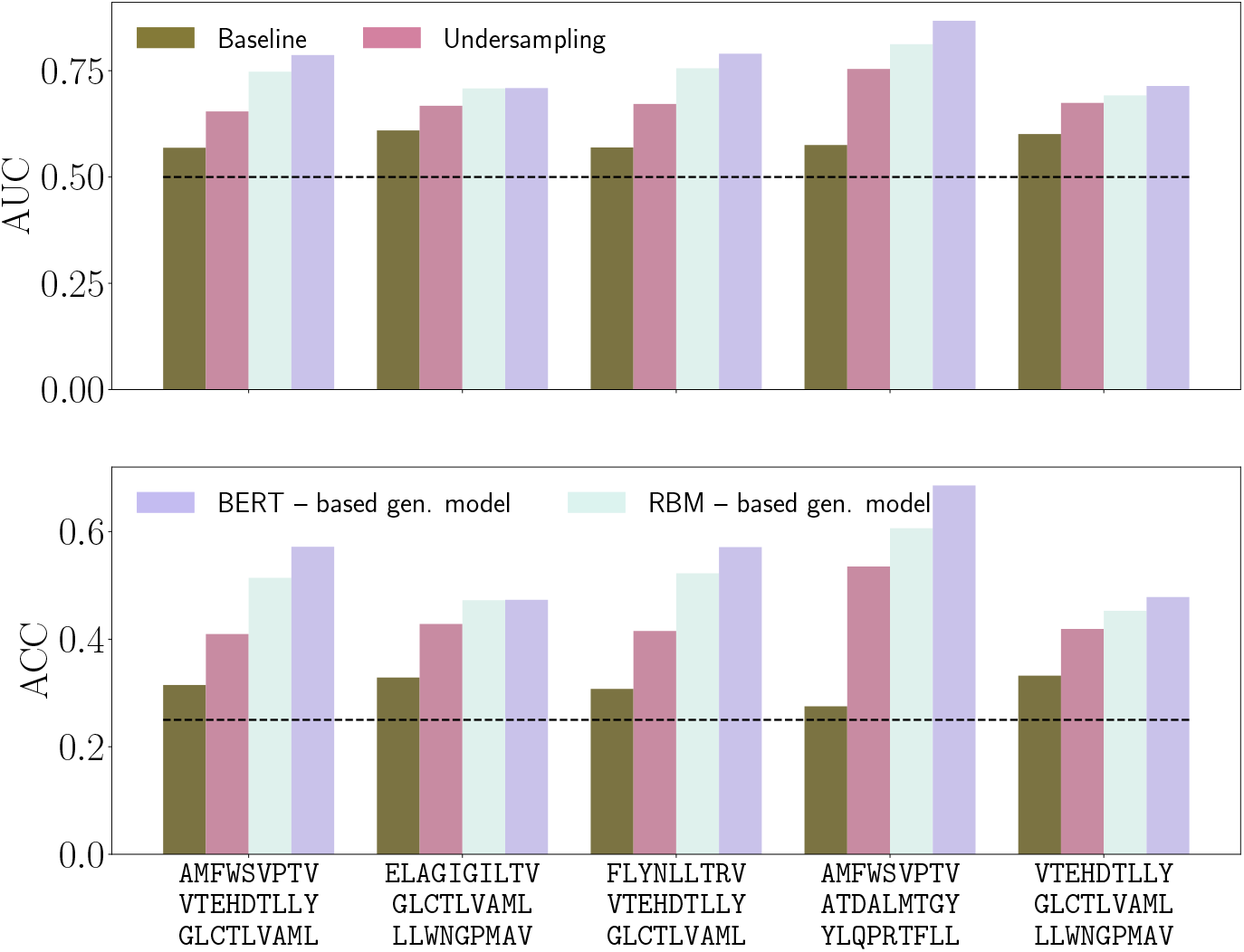
In-distribution performances of peptide-specific models. AUC and ACC scores of the predictive models for a multiclass classification task involving three peptide-specific sets of CDR3*β* and background CDR3*β* sequences, evaluated over a balanced test set of sequences of the same classes (in-distribution case). We compare performances for different training datasets, whose balance is restored through undersampling solely, or by generating new CDR3*β* sequences via an RBM- or BERT-based generative architecture. The baseline scores refer to an imbalanced training dataset, whose composition can be derived from the class sizes of each epitope as reported in App. 1; 250,000 background CDR3*β*s are used. Results confirm the benefit of both restoring balance in the training dataset and enlarging the peptide-specific CDR3*β* space through generative models. Dashed black lines indicate random performance levels.

Notice that, to obtain the increase of performances we observe in Fig. 3, it is crucial to balance the dataset with generative models powerful enough to capture non-trivial features in the distribution of the data, a task that both RBMs and BERT-like models are able to do; for more on this aspect, see App. 3.

Similarly with what we will report in the next section for pan-specific models, performances of the peptide-specific ones can also be assessed in an out-of-distribution setting, in which the test set includes sequences binding to unseen epitopes. As expected, model accuracy strongly depends on the similarity between the out-of-distribution and training data distributions, see App. 4.

### Pan-specific models for in-distribution predictions benefit from data augmentation

Pan-specific models, which take as inputs both the CDR3*β* and the peptide sequences, have gained interest, as they offer the possibility to identify binding patterns across different epitopes, and could potentially be used for binding predictions with rare or even novel peptides.

Based on the results from peptide-specific models, we now probe the benefit of restoring balance through the use of a pan-specific generative model, which we train over the full dataset from ***Grazioli et al. (2022)*** (see App. 1 for its composition), including abundant peptide-specific groups of CDR3*β*s. We then generate new peptide-specific sequences through Gibbs sampling initialized with natural sequences. The model takes as input peptide and CDR3*β* pairs of sequences and proposes random mutations across the sequence pair to sample new ones. We observe that only a small fraction (~ 3%) of generated data shows mutations across the peptide sequence: data augmentation overwhelmingly consists in the production of new CDR3*β* attached to the same peptides as in the training dataset.

An important hyperparameter is the size of peptide-associated groups after generation of new TCR sequences. In practice, we decide to set a threshold group size 𝒢 for data augmentation, above which peptide-specific classes are deemed as *sufficiently populated* and are thus not enlarged through the generative model. Groups having less than 𝒢 datapoints are all enlarged up to the threshold value. 𝒢 cannot be chosen arbitrarily large, as the generation of new data points is computationally expensive, and limited by the quality of training data and of the unsupervised model.

Once all positive data have been generated, we restore balance at class level by undersampling negative peptide-CDR3*β* pairs. Undersampling is randomly performed within each group, so that we are guaranteed that the dataset contains, on average, equal numbers of positive and negative pairs for each group, a requirement that we have observed to be important performance-wise.

We show in Fig. 4 that data generation leads to increase of performance for a vast majority of the peptides involved in the dataset. Even peptide-specific groups that have not been directly enlarged (red circles) can show a gain, arguably because the CNN architecture is leveraging the information of generated peptide and CDR3*β* pairs to transfer learning across all the different peptide groups. The scores also show that the generative model is particularly promising for heavily under represented groups (small triangles), with performance gains up to 20%.

**Figure 4.**
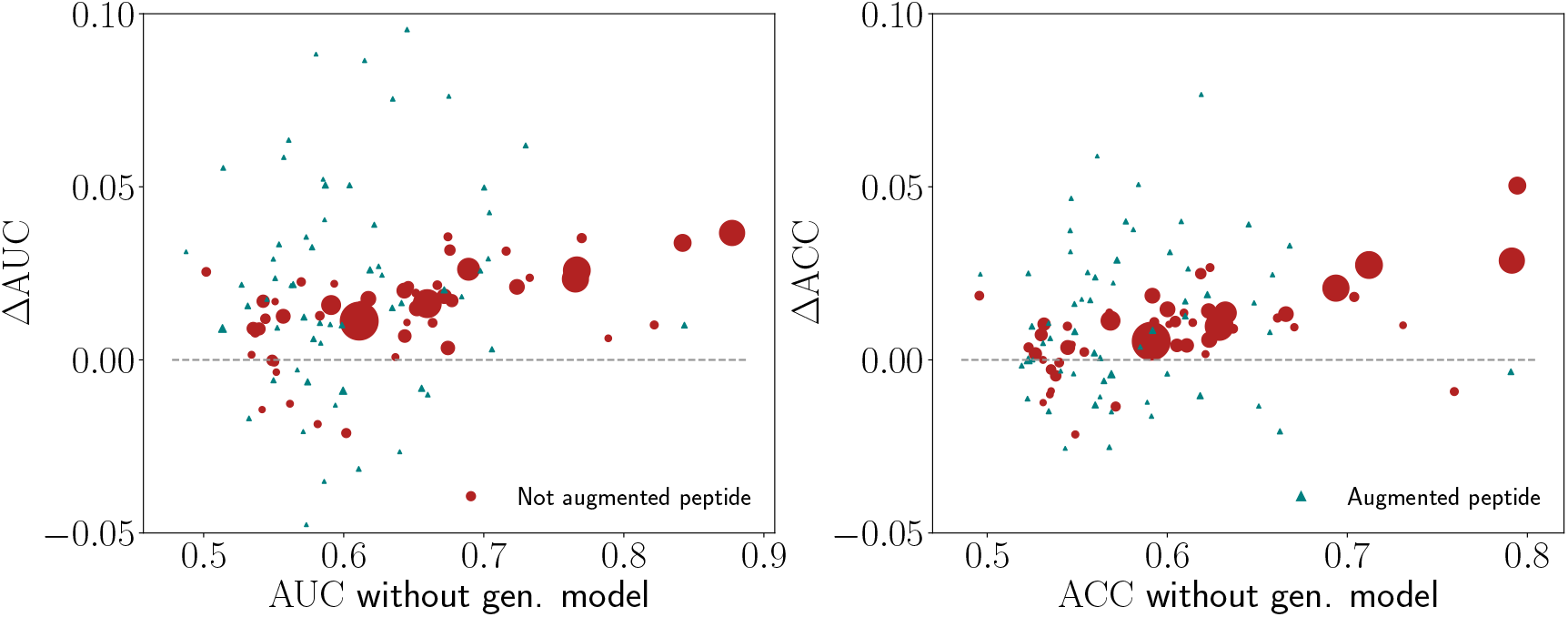
Pan-specific predictive model results. Differences in AUC and ACC scores when balance is achieved through undersampling of data only (no generation) or with data augmentation too. In the latter case, new CDR3*β* sequences were generated for under-represented groups of peptides only (triangular dots). Dot sizes are proportional to the raw group size of natural sequence pairs in the dataset. Here, 𝒢 = 400 (for more details on the choice of this threshold, see App. 5).

### Out-of-distribution performance on close enough sequences is positively impacted by data augmentation

Having established that our two-step pipeline is beneficial for in-distribution predictions, we set out to investigate whether this still holds for an out-of-distribution analysis. In this context, test data points are sampled from an external distribution – which may share only few features with the training data distributions. Consequently, out-of-distribution predictions can be very challenging. We expect performances to drop consistently, depending on how much the out-of-distribution data are distant from the training data in feature space.

In addition to predictions for peptide and CDR3*β* sequence data, we analyze out-of-distribution performances of our approach on synthetic data, which offer a realistic scenario where the ground truth is available and it is thus possible to quantitatively “measure” out-of-to in-distribution distances.

#### Binding predictions on unseen epitopes

We now ask whether our pan-specific model generalizes well on out-of-distribution TCR-peptide sequences, *i*.*e*. is able to capture the general properties underlying the binding process of receptors to antigens. As out-of-distribution test set we aggregate all the peptide-specific CDR3*β* sequences excluded from the training set during data pre-processing because they were strongly under represented (see Materials and methods); yet, we retain only groups that contains > 20 CDR3*β* sequences to have significant statistics. We report their abundances in Tab. 1.

Performances for this hard out-of-distribution setting are unsurprisingly worse than in the in-distribution case, in agreement with previous studies (***Croce et al., 2024)***. Nonetheless, we observe a correlation between the scores and the similarity between the unseen epitopes and the ones in the training data; in fact, clustering the CDR3*β* sequences based on the Levenshtein distance of their target epitope to the closest epitope in the dataset, we find that performances drop from AUC = 0.68 for close epitopes to random for distant epitopes (Levenshtein distance 11), respectively. We report the AUC/ACC metric results grouped by Levenshtein distance for all tested cases in App. 4.

**Table 1.**
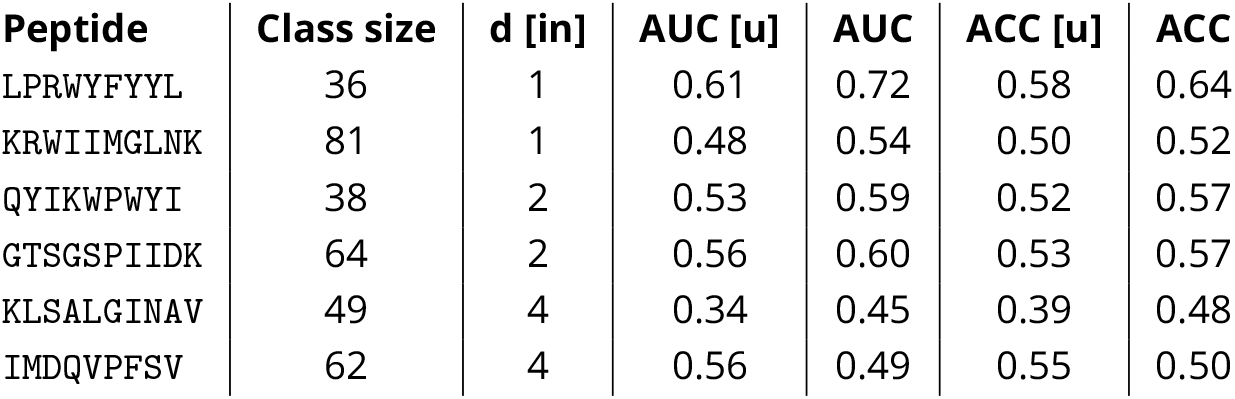
Out-of-distribution performances evaluated with AUC and ACC metrics across a test set composed of wild type binders to the target epitope and CDR3*β* sequences sampled from other unseen epitopes. For each prediction, we separately train a classifier with an enlarged training set containing also the synthetic binder of the target out-of-distribution epitope, generated through the pan-specific model (trained on the in-distribution dataset only, but with the target epitope given during the generative step, as explained in the text). The columns labeled with [u] refer to scores obtained balancing the training set by only undersampling the negative class, for comparison. The column d[in] represents the Levenshtein distance from the closest in-distribution epitope, showing that scores degrade when moving away from in-distribution data.

Improving out-of-distribution classification is crucial for the task of TCR-peptide binding predictions, as information about the binding properties of new antigens are rarely available. Here, we explore the possibility of predicting antigen binding towards unseen epitopes, exploiting the pan-specific generative model (*trained on the in-distribution dataset*) and *a partial knowledge of the out-of-distribution dataset* (the sequence of the new epitope). In practice, we use the generative architecture to sample CDR3*β*s binding to the target epitope; then, we include these data in the training set and test the performance on the natural out-of-distribution binder sequence data.

As a proof of concept, we evaluated this procedure on 8 different epitopes, whose natural CDR3*β* sequence have to be distinguished from other CDR3*β* with a different out-of-distribution specificity (see Tab. 1). We observe an improvement of performances for epitopes at small distance from their in-distribution counterparts.

Minimal model of out-of-distribution classification of TCR-peptide binding pairs To better understand how the properties of out-of-distribution data impact binding predictions, we repeat this analysis on controlled, synthetic data. We resort to dimeric lattice proteins (LPs) compounds, whose native folds are shown in Fig. 5a. LPs are synthetic proteins defined as self-avoiding paths over a 3 × 3 × 3 lattice cube, whose vertices carry the 27 amino acids. The two LP structures in the dimer represent, in order, the epitope and the TCR.

**Figure 5.**
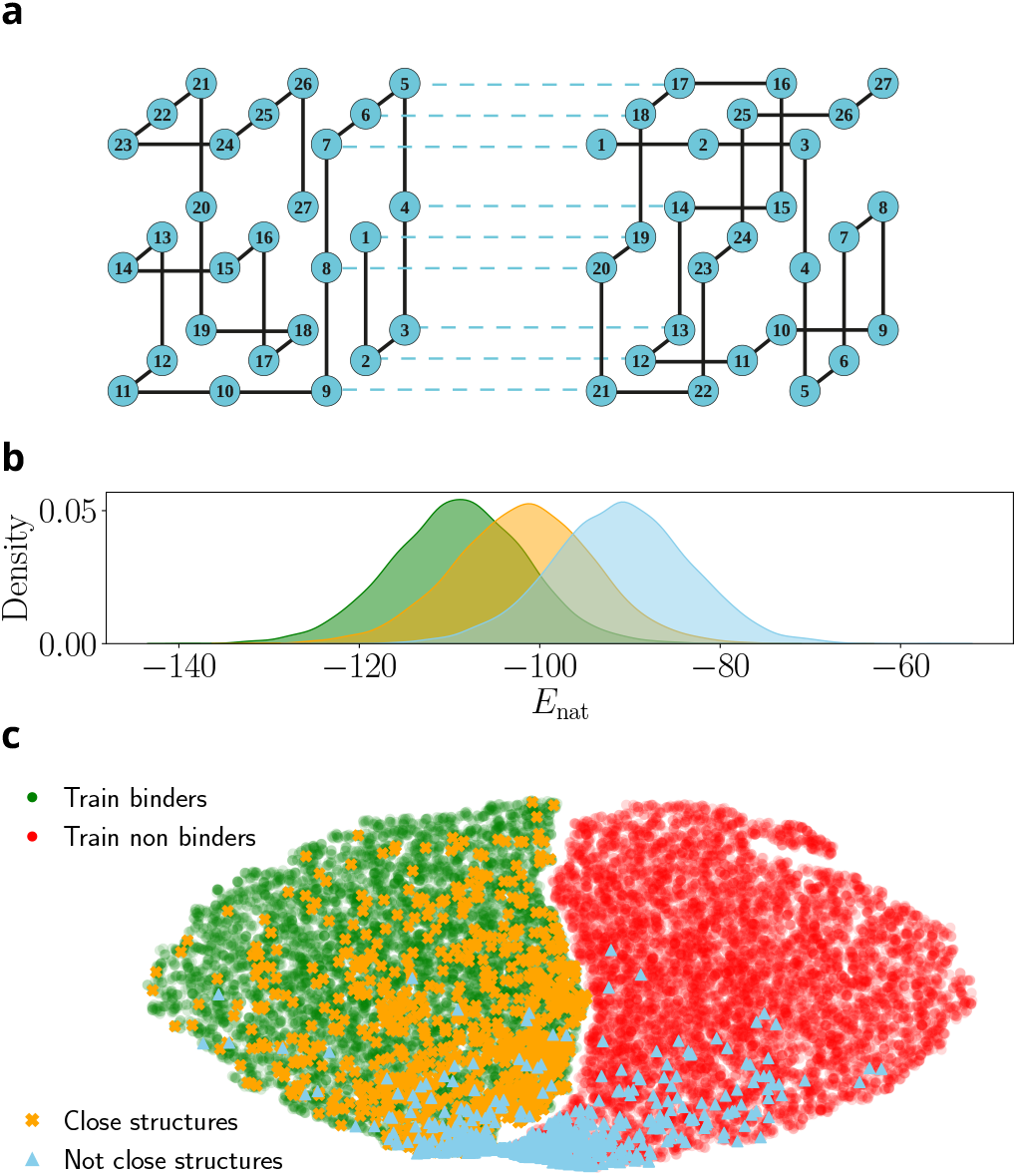
Out-of-distribution predictions on synthetic LP dimers for pan-specific case. **a)** The two structures of the proteins making the dimer, with amino acids defining a strong binding interaction, represented by the dotted lines. **b)** Histogram of single structure folding scores (the lower the better), computed according to the ground truth of the model (see App. 6, Eq. (16)). In practice we take MSA of binding sequences used for training and compute the folding scores in their native structure or as if they are in an out-of-distribution close structure or not close structure. The three distributions confirm the vicinity of the sequence data orange structures to the green ones, compared to blue ones. **c)** tSNE visualization of in-distribution training data (binder and non binders, green and red respectively) and out-of-distribution hold-out data (close and not close structures binders) over the embeddings of our CNN architecture. In this layer the classification is linear and we can see a clear decision boundary separating green and red data points. Accuracy on in-distribution data is ACC = 0.99. The close out-of-distribution data are similar to training data, hence the model performs well on these ones (ACC = 0.98); conversely, it gives poorer performances on the not close out-of-distribution (blue data points, ACC = 0.82).

Following ***Loffredo et al. (2023)***, we build dimeric LPs starting from single monomers running Monte Carlo (MC) evolution, and collect sequence data for multiple dimeric LPs. Spanning multiple dimeric structures, by changing the conformation of the self-avoiding paths on the lattice, allowed us to model different group specificities; details in App. 6. We collect MSAs of binding and non-binding pairs constituting the two classes in our dataset, and balanced both at class and group levels. A CNN classifier similar to the one introduced for natural data above is then trained over these data and reaches perfect classification in discriminating binding vs non-binding pairs – regardless of the group specificity, showing the simplicity of this in-distribution task.

To assess out-of-distribution performances we produce an additional dataset of binding pairs as follows. For each group of dimers in the training dataset, we collect sequence data corresponding to its closest dimer, *i*.*e*. with highest structural similarity. This dataset is referred to as *close out-of-distribution*, as we expect it to be very close to the training data in the feature space of the classifier. Similarly, we repeat the procedure with randomly picked dimers, which share few similarities with the dimeric structures defining the training data; we label such data as *not close out-of-distribution*. The structural similarity based on the ground truth folding scores is reported in Fig. 5b: for all in-distribution binding pairs sequences we plot scores for native, close and not close dimer structures; the orange distribution is effectively closer to the green one than the blue one, confirming a stronger structural similarity.

In Fig. 5c we report the tSNE 2d projections of the embeddings of the supervised architecture trained over the binary classification task for binding (green) versus non-binding (red) pairs of sequences, which allows us to visualize the neat decision boundary, giving very high ACC = 0.99. Within this feature space, on top of green and red points, we project out-of-distribution data points for the close and not-close cases. As expected, the latter is harder to classify as they share less features with the training data of the model (ACC = 0.98 and ACC = 0.82, respectively). Therefore, even in the framework of artificial data, we observe that out-of-distribution predictive performances depend on the degree of similarity with the in-distribution dataset.

## Discussion and conclusions

In this work we have introduced a framework that combines the use of unsupervised and supervised computational approaches to achieve reliable predictions of TCR specificity. Our pipeline relies on a two-step procedure, where we first leverage unsupervised network architectures to learn the probability distribution of peptide-specific CDR3*β* sequences or peptide-CDR3*β* binding pairs from which new informative sequence data are generated afterwards. Second, the generated data allow us to train supervised models for the final predictive task over balanced datasets, avoiding biases induced by class imbalance. We emphasize that restoring balance by augmenting the under-represented positive data class through generation yields better performances than downsampling the negative, over-represented class alone. The gain in performance is found not only for in-distribution data, but also for out-of-distribution tests. In this latter situation, the quality of predictions however decreases with the dissimilarity between the tested epitope and the ones present in the test set. This effect, and more generally the reasons why restoring balance improves classification performances, can be geometrically interpreted in the feature space of the classifier – see Introduction and App. 2.

The proposed pipeline resorts to unsupervised learning to balance the training set of a supervised classifier. A natural benchmark for performance is the direct use of the unsupervised model alone to classify the data. By fixing a threshold on the score that the model assigns to test sequences, measuring how likely they are sampled from the distribution of the positive data used for training, we may obtain predictions for class membership from the unsupervised model alone. This ‘unsupervised classification’ procedure generally yields worse generalization results than the full pipeline proposed here, as reported in App. 8. The decision boundary obtained by fixing a threshold on the score of the unsupervised model, which is trained on positive data only, does not coincides with the surface separating positive and negative features in sequence space, see App. 2.

Restoring balance in the data is crucial to improve predictive power not only when one designs peptide-specific models, a case in which the imbalance is present at class level only, but also for pan-specific models, for which the amount of data associated to each epitope may largely vary, with some peptide being heavily under–represented. An alternative approach could be that of combining peptide- and pan-specific models not only for the supervised predictive task – as already proposed in ***Jensen and Nielsen (2023)*** – but also for data generation; in this way, a limited number of generative models would be trained separately over the most abundant peptide-specific classes, while a single generative model would be trained over peptide and CDR3*β* pairs of sequences. Depending on the peptide, one of the two generative models could then be adopted. Furthermore, the effect of group imbalance could be reduced introducing a re-weighting factor for each group in the loss function of the generative model during training – inversely proportional to the group size.

The training data used in this study for peptide- and pan-specific models have been collected from publicly available databases, which curate and provide T cell receptor sequences and their cognate targets published in literature. Despite recent advantages and efforts to make available also negative assays of TCR-epitope bindings, such resources remain biased towards positive interactions since negative interactions are rarely reported in experiments. To obtain negatively/non– interacting pairs of sequences practitioners resort to various strategies, including the reshuffling of epitopes and CDR3*β* pairs. However, recent works have shown that the production of negative data may induce biases and impact the predictive power of models (***Dens et al., 2023; Ursu et al., 2024)***. Alternatively, negative data could be derived using pair complexes with low binding affinity generated through synthetic lattice-based receptor (***Akbar et al., 2022; Robert et al., 2022)***. Together with the increasing amount of available negative assays, we believe that these studies will contribute to better understand and limit the sources of biases stemming from negative samples within the context of TCR-epitope binding analysis.

In conclusion, our results demonstrate the benefit of reducing imbalance for both peptide– and pan–specific models, while suggesting to be more important when there is an heavy imbalance in the initial dataset of natural sequences. Preprocessing the dataset to restore balance is effective across multiple strategies, ranging from simple undersampling of abundant sequences to the generation of new peptide-specific CDR3*β* sequences with unsupervised architectures, including energy-based models and Transformers. The robustness of these gains suggests that even better TCR specificity predictions could be achieved with more powerful generative models and/or hyperparameter optimization. For instance, our learning pipeline could be easily extended by including the possibility to feed the CDR3*α* chain, VJ annotation or MHC class information, likely leading to performance improvement as shown in ***Sidhom et al. (2021)***; ***Montemurro et al. (2021)***. Similarly, better performances could be reached by simultaneously training the generative and the classifier models, rather than one after the other. In addition, to study the impact of the training set composition on the performances alone, we did not finetune any hyper parameter and we trained all our models with the same number of total iterations over the batch, *i*.*e*. by rescaling the epochs according to the training set size.

To end with, let us emphasize that our generative models allow for the possibility of producing new, putative CDR3*β* sequences with desired binding specificity that could be tested experimentally. Thanks to its methodological simplicity and flexibility, our learning framework is not limited to the context of T cell receptors specificity predictions, and could be applied to other sequence– based computational problems where heavy imbalance impedes the proper training of predictive models (*e*.*g*. ***Haque et al. (2014)***; ***Ming et al. (2023)***; ***Rana et al. (2022)***; ***Li et al. (2022)***).

## Materials and methods

### Generative models for data augmentation

Generally speaking, unsupervised machine learning aims to learn an energy landscape by inferring parameters of a user-defined probabilistic model *P*_model_ over the data presented (in our case, the peptide specific CDR3*β* sequences, or the peptide + CDR3*β* binding sequences). The model is trained (or fine-tuned) separately over each class and is then asked to generate new sequences compatible with real ones by sampling from the learned probability landscape. To generate new sequences we make use of Restricted Boltzmann Machines (RBMs) (***Tubiana et al., 2019; Bravi et al., 2023)*** and of Large Language Models (LLMs), specifically we use architectures based on Bidirectional Encoder Representations from Transformer (BERT) adapted to handle CDR3*β* sequences (***Wu et al., 2024)***.

#### Restricted Boltzmann Machines

RBMs are bipartite graphical models including a set of *L* visible units **v** = (*v*_1_, ⋯, *v*_*L*_) and *M* hidden (or latent) units **z** = (*z*_1_, ⋯, *z*_*M*_). Only connections between visible and latent units are allowed through the interaction weights 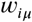. RBMs define a joint probability distribution over **v** and **z** as the Gibbs distribution

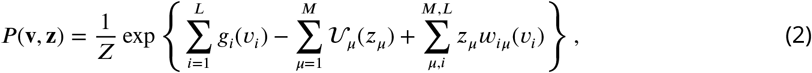

The joint probability above is specified by a set of parameters whose values are inferred from the data (here the biological sequences). They consist of i) the set of single-site biases 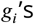 acting on visible units that capture the amino acid usage at each sequence position; ii) the potentials 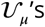 acting on hidden units, here assumed to have dReLU shape (***Tubiana et al., 2019)***, and iii) the set of weights 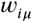 coupling the hidden and visible layers.

The probability of data *P* (**v**) is defined as the marginal of the joint probability over the data itself

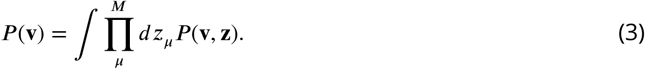

All the parameters in the model are learned by maximizing the marginal log-likelihood

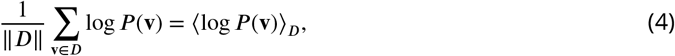

where *D* is our dataset and ‖ *D* ‖ its size. Due to the nature of RBMs, all data needs to have the same dimension to be accepted: in practice, we align the CDR3*β* receptors before feeding them into the model.

From the definition of the model reported in Eq. (2), let us define the RBM energy as

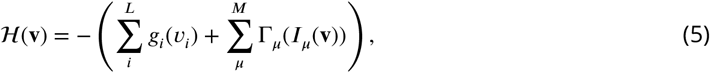

where we introduced the shorthand notation 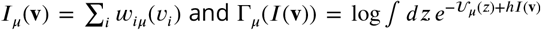. For a generic parameter 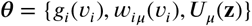, the rule to infer its value is given by the gradientascent equations stemming from log-likelihood maximization,

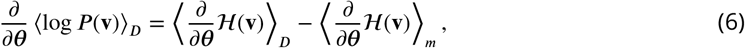

where ⟨⋅⟩_*m*_ stands for the average over the model, ⟨*u*(**v**) _*m*_⟩ = Σ_v_ *P* (**v**)*u*(**v**). In addition, regularization terms can be added to control the values of inferred parameters, by enforcing sparsity on the weights through *L*_1_-penalty to avoid overfitting and by controlling their norm through 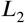-penalty to prevent divergences. In practice we resort to a 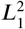 regularization scheme that consists in adding to the log-likelihood of data the following penalty term

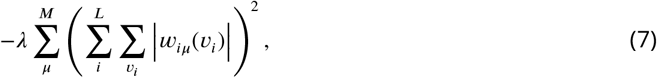

where λ sets the regularization strength. It has been suggested that such regularization also helps to improve the generative properties of RBMs. The parameters used for learning of peptide-specific CDR3*β* distributions are:

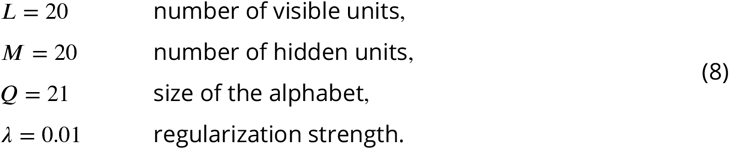

Once the full set of parameters has been inferred from the training data, we can sample from the probability distribution *P* (**v**) to obtain new data (in our case, new peptide specific CDR3*β* receptors). Here we sample using Alternate Gibbs Sampling (AGS), which consists in alternatively sampling from the RBM’s visible layer while keeping the hidden layer fixed from *P* (**v**|**z**) and vice versa from *P* (**z**|**v**). The Monte Carlo Markov Chain (MCMC) is initialized starting from sequence data in the dataset and new sequences are collected after some steps of thermalization to avoid sampling data correlated with real ones.

### BERT-based architectures

We consider a generative model of amino acid sequences based on the Bidirectional Encoder Representations from Transformer model architecture (***Devlin et al., 2019)***. Transformers models are build through a series of attention blocks and feed-forward layers, whose aim is to capture interactions across the input sequence through learning how strongly the embedding of each token in the input is affected by the other tokens (the context). Stacking multiple attention blocks allow the model to learn complex structures within the embedding of the input data, such as semantic, causal and grammatical properties of the data.

The model takes as input for training CDR3*β* sequences for the peptide-specific case and pairs of peptide and CDR3*β* sequences for the pan-specific case; the sequences are formatted via the tokenizer retrieved from ***Wu et al. (2024)***, which contains a total of 26 tokens spanning the 20 amino acids and some special tokens, such as the prefix and suffix token and the masking one. An additional token & is included for the pan-specific case to model the peptide – CDR3*β* separation in input data (*e*.*g*. `GILGFVLT` & CASSLDGTVQYF). Sequences do not need to be aligned, and the inputs are padded with an attention mask. The input tokens are thus padded through the embedding layer to get a continuous representation vector of the data: such embedded vector (together with a positional encoding vector that retains the positional information of amino acids in the input sequence) goes through the model attention blocks. The output feature vector leaves in a *L* × 768 dimensional space, where *L* is the input length, and is fed to a task-specific head for masked language modelling.

The training over our dataset is carried out performing masked language modelling (MLM) objective, where we mask a fraction of input token in the dataset and train the architecture to predict the correct ones. This allows the model to learn the grammar underlying the CDR3*β* and peptide sequence patterns. For the peptide-specific case, the generative model is obtained by fine-tuning the TCR-BERT model from ***Wu et al. (2024)*** through few epochs of MLM objective over the peptide-specific CDR3*β* data; this leverages the transfer-learning ability of the model that has trained over unlabelled TCR sequences to capture distinctive features of the peptide-specific CDR3*β* sequences. For the pan-specific model, we train the model from BERT weights over the full training dataset; at variance with BERT model, we set the maximal positional embbeding length allowed to 64, due to the shortness of sequence data compared to text data.

For the (fine-)tuning procedure, given a TCR dataset ℳ and a masking pattern 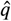, we minimize the following training loss

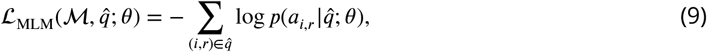

where *θ* is the set of all parameters in the model and (*i, r*) run over the residue position and sequences, respectively. The conditional probabilities *p* for each (*i, r*) are computed using the softmax-normalized model output values - the *logits* - for each symbol in the alphabet. For each input sequence, we mask 15% of amino acids as done in the original training ***Devlin et al. (2019)***. All hyperparameters during fine-tuning are the same as the ones used for training in ***Devlin et al. (2019)***.

Once the model has been (fine-)tuned on the MLM downstream task, we leverage its generative power following the iterative masking scheme proposed in ***Wang and Cho (2019)***; ***Sgarbossa et al. (2023)*** for protein sequences. In practice, each step of this procedure works as follows:

1. we randomly mask with probability *q* each entry of the CDR3*β* sequence or we leave it unchanged with probability 1 − *q*. This defines a masking pattern 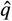;
2. we feed the masked sequence to the model and replace masked entries sampling from the softmax (normalized) distribution of the model logits at inverse temperature *β*;
3. we repeat steps (i) and (ii) for *T* times and then we store a new sample sequence.

In practice, we set *q* = 0.2, *β* = 1 and *T* = 20 and we repeat this scheme many times starting from the last configuration to obtain the desired number of TCR sequences. At the end we merge all the generated CDR3*β* and eventually drop duplicates: the remaining samples constitute the new set of sequences that will augment peptide specific sets.

Note that our protocol does not allow insertions or deletions across the sequence, as only residue tokens can be masked: in this way, the length distribution of training and generated data remains the same.

### Model architecture for supervised predictions

We then want to train a classifier from the training data (CDR3*β* sequences, or CDR3*β*-peptipe pairs, with the corresponding class labels). Many architectures, of widely varying complexity, were implemented in the context of TCR specificity predictions. Hereafter, we consider one architecture, and expect our results on restoring balance through generative models of CDR3*β* to hold for other network architecture chosen for the supervised step (***Loffredo et al., 2024)***.

In practice, we implemented a 1-D Convolutional Neural Network (CNN) to make binary and multiclass predictions on TCRs specificity. Our architecture takes as input raw sequences and returns a single value corresponding to the target class label assigned by the model (*e*.*g*. if the CDR3*β* considered binds one of the epitopes or it belongs to the bulk repertoires). CDR3*β* sequences are first padded to a max length *L* = 30, then translated into score matrices using the BLOSUM50 encoding matrix with no gap or special amino acid included, see ***Henikoff and Henikoff (1992)***. Hence, each sequence is mapped into a score matrix of size *L* × 20. The encoded input is then fed into the network architecture, consisting of five convolutional layers having 16 filters and different kernel sizes {1, 3, 5, 7, 9}. The resulting feature vectors go through a batch-normalization layer to reduce overfitting and are then concatenated to pass through a dense layer with 16 hidden neurons. An additional batch-normalization layer is applied and the resulting 16 dimensional feature vector is used to visualize the embeddings constructed by the model starting from raw data. The final classification is performed over such embeddings by feeding the feature vector to a layer with a number of neurons equal to the number of classes in the dataset and having softmax activation function (for binary classification tasks, we actually use one neuron and sigmoid activation function). We use the ReLU activation function throughout the network. Adam optimizer with learning rate *η* = 0.001 and categorical (or binary) cross-entropy loss function are used for learning with a batch size of 128 samples.

Performances are evaluated using accuracy (ACC) and Area Under the receiver operating characteristic Curve (AUC) metrics on balanced test sets, unless otherwise specified. The former is defined as

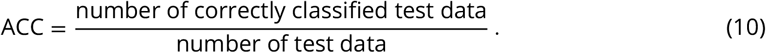

In the case of binary classification, the receiver operating characteristic (ROC) curve is defined as the parametric curve in the False Positive Rate–True Positive Rate plane as a function of the threshold used to assign the binary label to the output of the sigmoid activation function of the readout layer. In the case of multiclass classification, AUC is defined as a uniform average of the binary AUCs of all possible combinations of classes (one-vs-one):

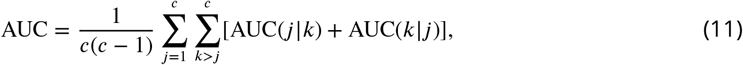

*c* being the number of classes.

## Datasets collection

Supervised and unsupervised models developed in this work have been scored using multiple type of synthetic and real data. In all cases – after preprocessing – positive and negative data were merged together in a final dataset with a proportion *ρ*_*P*_ :*ρ*_*N*_ that can be tuned. The dataset was then split into three parts for training, validation and test. Subsampling from the previous test set, we also construct a balanced test set having the same numbers of positive and negative examples. We refer to performance on the latter set as in-distribution performance. We then assess model predictions on unseen (balanced) examples to define out-of-distribution performance.

We provide below details on TCR-peptide binding data we used to analyze peptide- and panspecific models with our pipeline.

### TCR-peptide data for peptide-specific models

Sequence data for TCR-peptide specificity predictions were retrieved from the Immune Epitope Database (IEDB) (***Vita et al., 2018)*** as of June 2023, filtering the Database entries as follows. We focus on human host immune response and set the pMHC restriction to the HLA-A*02:01 complex, limiting the peptide length to 8 − 11 amino acids. Paired CDR3*β*-epitope sequences were thus identified as those for which a T cell assay was reported ‘Positive’ or ‘Positive-High’ and never ‘Negative’. We collected in this way sequences of CDR3*β* with specificity to a set of different epitopes, listed in Tab. 2. A summary of the resulting dataset, with the abundances of CDR3*β* sequences for each peptide, can be found in App. 1.

**Table 2.** List of the epitopes selected to collect CDR3*β* specific sequences in order to form the database used in the analysis of peptide specific models in Results.

| Epitope | Sequence |
| --- | --- |
| TKT-R438W from Melanoma cancer | AMFWSVPTV |
| NS3<sub>1436-1444</sub> from Hepatitis C virus HCV | ATDALMTGY |
| MART1<sub>27-35</sub> from Melanoma cancer | ELAGIGILTV |
| SEC24A from Melanoma cancer | FLYNLLTRV |
| BMLF1<sub>280-288</sub> from Epstein-Barr virus | GLCTLVAML |
| peptide from Yellow Fever virus | LLWNGPMAV |
| pp50 from human cytomegalovirus CMV | VTEHDTLLY |
| peptide from Spike protein S269 of SARS-CoV-2 | YLQPRTFLL |

Background sequences of CDR3*β* considered as those non-self reactive are taken from the database assembled by ***Isacchini et al. (2021)***, which merges together (i) unique clones from the 743 donors of the cohort in ***Emerson et al. (2017)*** and (ii) the dataset of CDR3*β* sequences from healthy donors in ***Dean et al. (2015)***. The full dataset is then sub-sampled at random for computational purpose and we retain 10^6^ background sequences.

Whenever sequence are fed into the RBM learning, they are required to have the same input length. Therefore, CDR3*β* sequences are aligned with an Hidden Markov Model (HMM) using as alignment profile the one built in ***Bravi et al. (2021)***. Aligned CDR3*β* sequences have fixed length of 20 amino acids. To train the supervised model with RBM-generated sequences we drop all the gaps inserted by the alignment procedure, as our deep classifier can handle sequences of different lengths.

As anticipated in the introduction, the assembled dataset suffers from imbalance between peptide-specific CDR3*β*s and the large abundance of bulk unlabelled data. In addition, some epitopes (*e*.*g*. GLCTLVAML, `LLWNGPMAV`, `YLQPRTFLL`) have thousands of known CDR3*β* binders, while others have hundreds of positively tested CDR3*β*s only, which results in further imbalance between the epitope-associated classes.

To avoid misleading predictions due to imbalance biases, we balance the training set in such a way that all classes are equally represented (Fig. 2). To evaluate performances in a fair way, we consider different metrics over a balanced test set containing all classes in the same proportion in order to factor out any source of bias. We use ACC and AUC metrics to assess our performances (see Model architecture for supervised predictions).

### TCR-peptide data for pan-specific models

The dataset used to train and test pan-specific models is taken from ***Grazioli et al. (2022)***, and contains both positive and negative interacting pairs of CDR3*β* and peptide sequences spanning 118 different epitopes. Natural data are taken from publicly available sources, namely IEDB, VDJdb, McPAS-TCR, MIRA (***Klinger et al., 2015; Nolan et al., 2025)***, and from 10X Genomics assays. Among them, we only retain peptides with at least 130 positive binder CDR3*β* representatives to obtain a sufficiently large dataset for training both the generative and the classifier models. This results in a total of 126,882 experimentally tested pairs of interacting CDR3*β* and peptide sequences, and a collection of 405,176 non-interacting sequence pairs. These negative examples consist for a minor part of experimentally assessed ones, while a large part of them is obtained by random mismatching of CDR3*β* and peptide sequences from the positive class; we further enlarge the negative set by pairing peptides with CDR3*β*s randomly chosen from the previous bulk repertoire. A summary of this dataset, with the absolute and relative abundances of CDR3*β* sequences for each peptide, can be found in App. 1.

The resulting dataset is plagued with two sources of imbalance:

1. positively interacting pairs are strongly under-represented compared to negatively interacting pairs – we refer to this as *class-level* imbalance;
2. within the positive class, few peptides are strongly over-represented compared to others – we refer to this as *group-level* imbalance (***Sagawa et al., 2019; Idrissi et al., 2022)***.

Notice that there is possibly group imbalance within the negative class based on the different ways to assemble negative sequence pairs through mismatches. We qualitatively observe that this imbalance has minor effects on performances and ignore it in the following.

The group imbalance could, in principle, also hinder the generative process of new peptide-specific CDR3*β* binding pairs for pan-specific models. In fact, a pan-specific generative model trained on peptide and CDR3*β* sequences pairs might effectively learn only the most-represented instances of epitopes, i.e. the most abundant groups in the positive class, making our pipeline ineffective. Comparison of this generative approach with multiple peptide-specific generative models (one for each epitope) shows that the impact of group imbalance is negligible, see App. 7.

## Code and data availability

The code to reproduce the analysis in this paper can be found at https://github.com/Eloffredo/RestoringTCR. Sequence data for TCR-peptide predictions are public and were retrieved from the Immune Epitope Database (IEDB) ***Vita et al. (2018)*** as of June 2023 and from TChard ***Grazioli et al. (2022)***, available at https://zenodo.org/records/6962043; utilities to collect them can be found in the above GitHub repository.

### Author contributions statement

E.L., M.P., S.C., R.M., designed research; E.L., M.P., S.C., R.M. performed research; E.L., M.P. analyzed data; E.L., M.P., S.C., R.M. wrote the paper

## Acknowledgments

We acknowledge funding from the CNRS - University of Tokyo “80 Prime” Joint Research Program and from the Agence Nationale de la Recherche (ANR-19 Decrypted CE30-0021-01 to S.C. and R.M.). E.L. thanks Andrea Di Gioacchino for interesting suggestions during the early stages of this work and his help with the use of the alignment software of receptor sequences. M.P. thanks Riccardo Capelli for discussions. The authors thank Victor Greiff, Eugen Ursu and Aygul Minnegalieva for comments and for the careful reading of the manuscript.

## Appendix 1

### Composition of the datasets used in the analysis of the main text

We report in Tab. 1, 2 the composition of the datasets used in the study of peptide- and pan-specific models in Results. Both were collected following Datasets collection.

**Appendix 1—table 1.** List of peptide-specific classes in the dataset used in the analysis of peptide-specific models. We report sizes of the peptide-specific classes defined by peptide sequences in the TCR dataset we used to study peptide-specific models in Results. Imbalance ratios for the 5 cases reported in Fig. 3 can be obtained from the relative abundances of the corresponding triplets of classes.

| Peptide | Class size | Peptide | Class size |
| --- | --- | --- | --- |
| AMFWSVPTV | 99 | ATDALMTGY | 160 |
| ELAGIGILTV | 821 | FLYNLLTRV | 142 |
| GLCTLVAML | 6364 | LLWNGPMAV | 1872 |
| VTEHDTLLY | 200 | YLQPRTFLL | 687 |

**Appendix 1—table 2.**
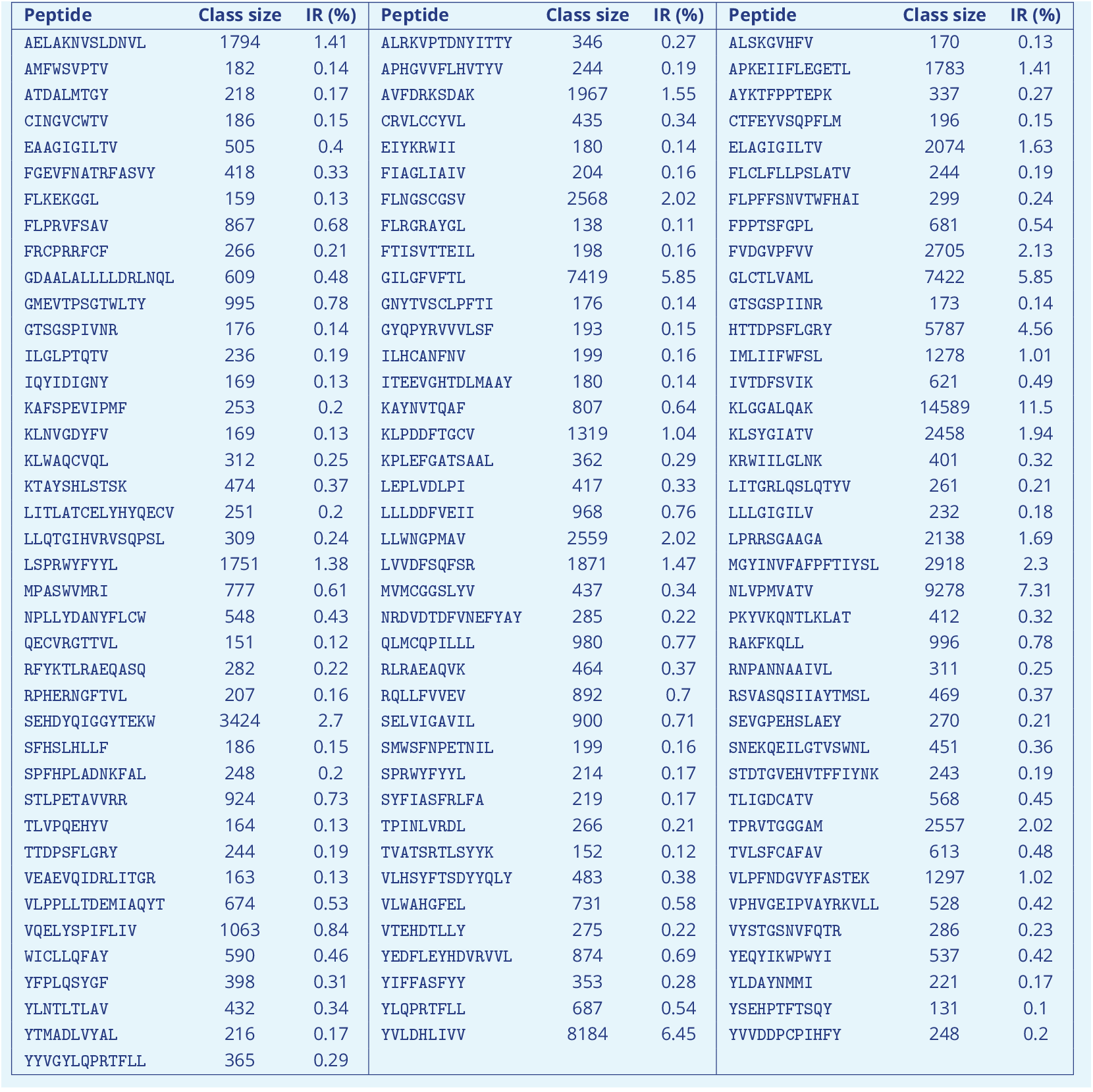
List of peptide-specific classes in the dataset used in the analysis of pan-specific models. We report sizes and imbalance ratio details on the peptide-specific classes defined by peptide sequences in the TCR dataset we used to study pan-specific models in Results. Only few peptide-specific classes have more than 1% of CDR3*β* binding sequences, causing heavily imbalance. Details refer to the dataset before training/test splitting.

## Appendix 2

### Geometrical interpretation of restoring balance

Supervised architectures solve the classification problem through the learning of a decision boundary in the high dimensional embedded feature space of input data. In this regard, achieving good performances means finding a good feature map where the input data points are well clustered based on their class identities. How does our restoring balance procedure via generative models of sequences affect the supervised learning process? We aim to visualize such effect to seek for a geometrical interpretation of re-balancing data points among different classes.

In particular, our supervised architectures used for both peptide- and pan-specific analyses employ a fully connected read out layer at the end of multiple convolutional layers. Thus, the model applies a series of transformations to map the input dataset into an higher dimensional space, which is passed to a fully connected layer where an hyperplane or a set of intersecting hyperplanes are found to perform binary or multi-class classification, respectively. Thanks to their simplicity, models with a single dense layer like Support Vector Machines (SVMs) allow for a mathematical formulation and some predictions on the model performance can be derived at theoretical level; within this framework, also the issue of learning under imbalanced datasets can be studied. For example, in ***Chaudhuri et al. (2023)*** the authors showed on common imbalanced benchmark datasets that undersampling helps classification as opposed to imbalanced learning; also, in ***Loffredo et al. (2024)*** we characterized the performances of an SVM under imbalance and studied the benefit of restoring data balance via under- and oversampling on synthetic data, showing that augmenting the under represented classes yields best performances. The key finding of such studies – that focus on binary classification – proves that learning under imbalance shifts the optimal decision boundary towards the under represented class and tilts it away from the optimal direction (see Fig. 1a, main text).

To visualize this effect, we consider a simple 1-hidden-layer network with linear activation and sigmoidal output, which we refer to as 1-Dense Network (1DN), and the CNN model considered above. In the two cases, the input-output map is given by

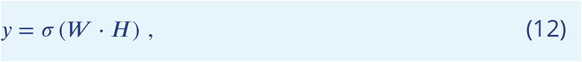

where *W* are the weights of the fully connected readout layer and the feature vector *H* is given by *H* = *W* ^(0)^*X* (1DN) or *H* = *f*_CNN_(*X*) (CNN). We train both our supervised CNN model and the 1DN model with a binary classification task (`LLWNGPMAV`-specific and bulk CDR3*β*s) using hinge loss. After training on the same dataset, we feedforward the test datapoints in the two models: we call *H*_±_ the two class centers of the test set,

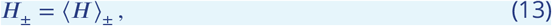

where the mean is performed over the two test set classes. The weight vector represents the direction of the decision boundary and should be aligned to the distance vector connecting the classes, *H*_+_ − *H*_−_; we quantify this alignment computing the normalized dot product

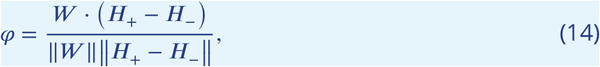

for both models, *φ*^CNN^ and *φ*^1DN^. Our hypotheses on the geometrical effect of learning with imbalance implies that we should find a value of *φ* closer to 1 when the model is trained over balanced datasets. Thus we run experiments on different training set compositions, tuning the fraction of negative examples (bulk CDR3*β*s) in the range [50%, 75%], averaging the quantity *φ* over 50 trials. Results on the simple 1DN and the deep CNN are in agreement, with the dot product dropping from *φ*^1DN^ = 0.87 (ACC = 0.62) to *φ*^1DN^ = 0.78 (ACC = 0.5) for the 1DN and from *φ*^CNN^ = 0.73 (ACC = 0.66) to *φ*^CNN^ = 0.62 (ACC = 0.5) for the CNN.

## Appendix 3

### On the choice of the generative model

Our learning framework is based on the idea that restoring balance through a combination of undersampling the strongly over-represented classes and augmenting under-represented classes via generative models. We believe such approach can yield better performances as the supervised network is provided with new informative peptide-specific CDR3*β* sequences; yet, the quality of generated sequences is crucial to prevent a loss of information. To discuss this point, we benchmark our specificity prediction performances with a simple model that requires zero training: a profile model (PM) that generates new CDR3*β* sequences by independently sampling each site of the sequence based on the peptide-specific class sequence profile (see Fig. 1). Since this model captures only first-order statistics of the sequences we expect it to generate less informative CDR3*β*s and thus we end with poor specificity predictions (see Tab. 1 for the scores obtained using this method for the cases reported in Fig. 3 of the main text).

The quality of data generation is also based on the stringency of the Gibbs sampling procedure, *i*.*e*. how easily we allow the generative process to accept random mutations. The degree of randomness in the sampling scheme is set by the temperature value *β* in the softmax outputs of the model: rescaling the model logits by *β*, low values of *β* flatten the softmax output distribution so that amino acids mutations are randomly accepted regardless from the underlying CDR3*β* distribution learned. To visualize this effect, we take a peptide-specific classification task and restore balance by generating random CDR3*β* sequences (see Fig. 2); at some point (*ρ*_rand._ > 50%) the random sequences completely take over the CDR3*β* natural sequences and make the specificity prediction problem impossible to solve, yielding random predictions.

**Appendix 3—figure 1.**
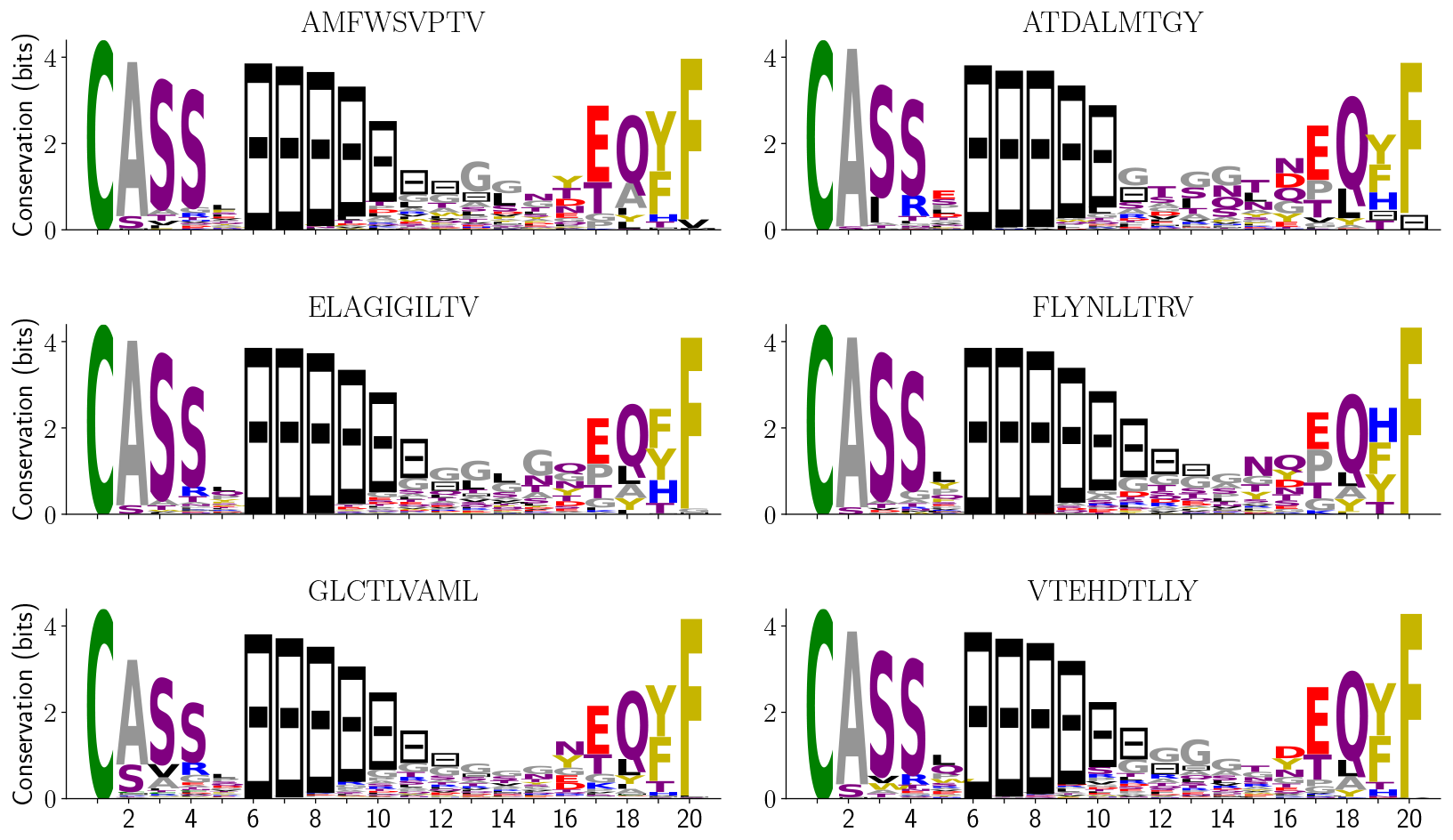
Sequence logos of peptide-specific CDR3*β* classes. We report the sequence logos profile for peptide-specific classes of CDR3*β* sequences, after alignment to maximal length *L* = 20. The PM is learned over such profiles for each class. Sequence logos show high conservation of the CASS motif and of the last amino acid, while there is more variability in the central region which is indeed responsible for the binding affinity.

**Appendix 3—figure 2.**
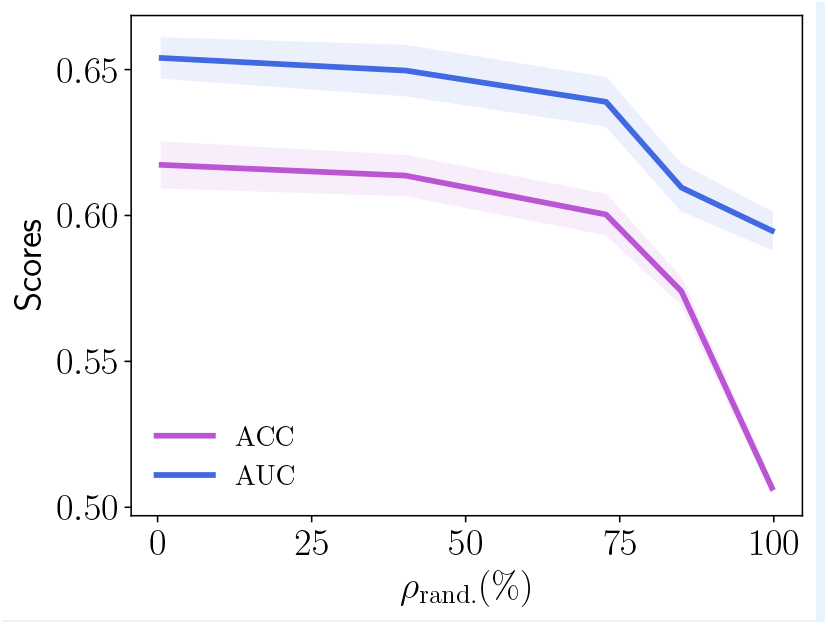
Performance drop due to randomness effect. Here we report AUC and ACC scores for a `VTEHDTLLY`-specific model that has been trained over a dataset containing a fraction of random sequences *ρ*_rand._. When the generative model is not good enough and is adding noise to the under represented class of data, we can observe the performances drop down to a random classifier.

**Appendix 3—table 1.**
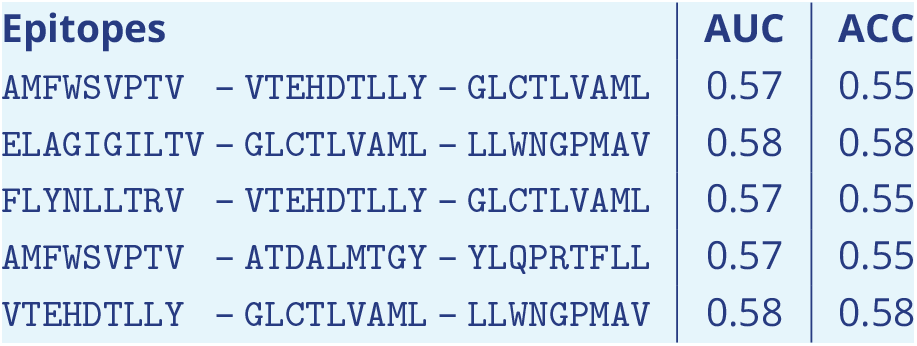
Performances scores for peptide-specific models with PM generative model. The supervised architecture is the same and it is trained as in Fig. 3 of the main text, on the same datasets with balance restored via generation of new CDR3*β* samples.

## Appendix 4

### More on out-of-distribution tasks

#### Performances of peptide-specific models strongly depend on the choice of the out-of-distribution data

Similarly to what done for the pan-specific framework in Results, we study out-of-distribution performances for single-epitope TCR specificity. Our peptide-specific model has learned the key properties of CDR3*β* binding to a selected epitope as reported in the main text, and it remains to be seen to what extent such features can be transferred to predict binding towards other target epitopes against bulk repertoires. We start by considering synthetic data.

##### Lattice proteins

We take data from ***Loffredo et al. (2023)***, where the authors built dimeric LPs starting from single monomers running Monte Carlo (MC) evolution at variable intra-dimer interaction strengths (see App. 6 for more details). We collect sequence data in the form of Multiple Sequence Alignments (MSA) at three steps during MC evolution, namely at beginning, intermediate and endpoint of evolution: the MSAs will constitute non binder, weak binder and strong binder data, respectively. We train a supervised model over strong and weak binder classes and use non binders as out-of-distribution data, which are thus closer to weak binders than to strong ones. We plot in Fig. 1a the binding probability distributions for these three classes of binders.

**Appendix 4—figure 1.**
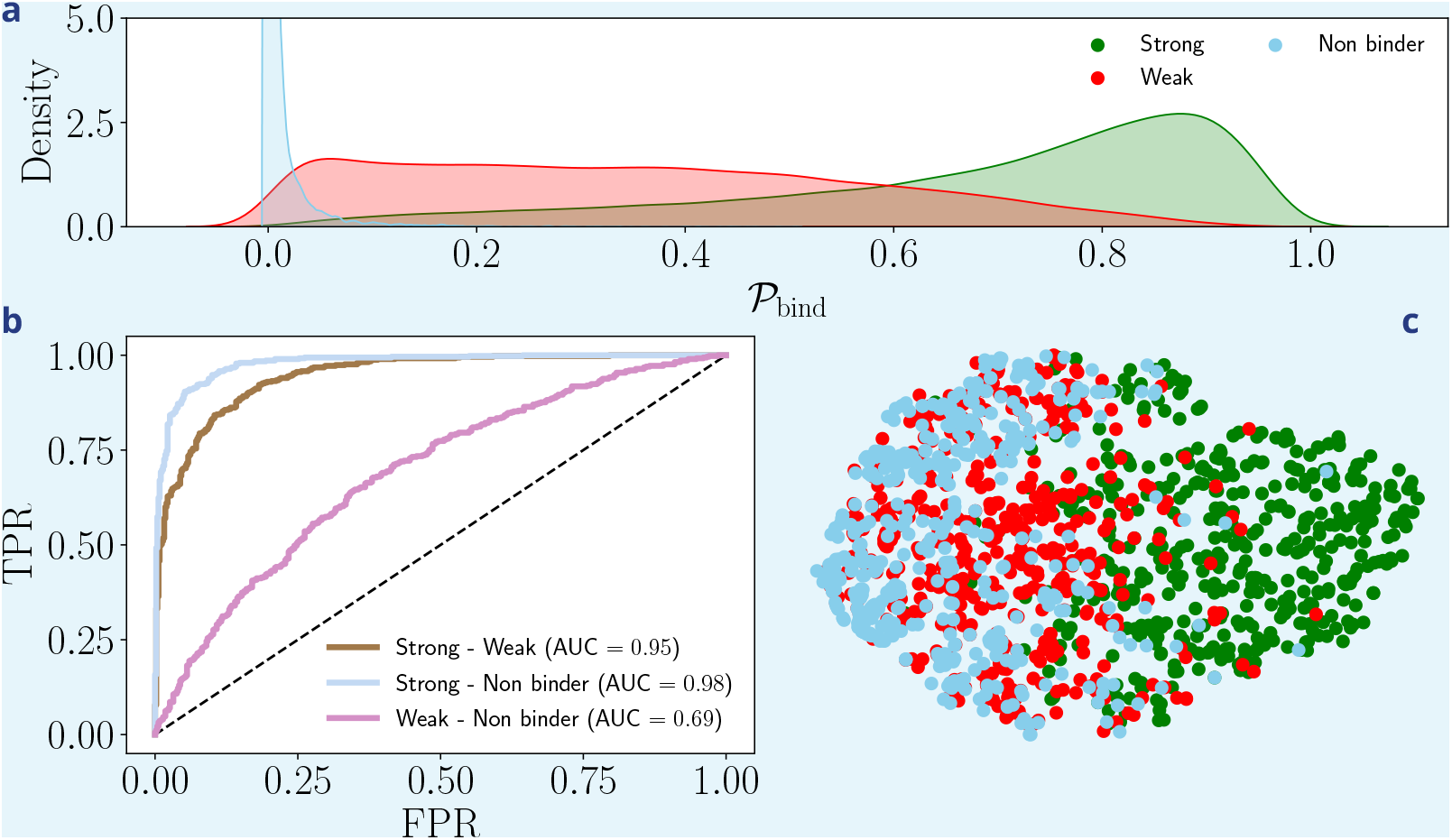
Out-of-distribution analysis on synthetic Lattice-Protein dimers. **a)** Densities of binding scores 𝒫 _bind_ for strong, weak and non-binding compounds. The y-axis is cut for visualization purpose, as “Non binder” compounds concentrates around zero score. **b)** Receiver Operating Characteristic (ROC) curves for in-distribution (Strong - Weak) and out-of-distribution (Strong - Non binder and Weak - Non binder) test sets. The classification weak-vs-non binders has worst performances. **c)** tSNE visualization of the embeddings (last feature layer) produced by our CNN architecture for in-distribution test data (strong and weak binders) and out-of-distribution hold-out data (non binders). Classification is carried out from linear combinations of the embeddings of the input data points; better separation of clusters reflects higher model performances.

The trained classifier achieve great performances on the in-distribution test set for strong-vs-weak binders classification (AUC = 0.95) and do even better for the out-of-distribution strong-vs-non binder classification (AUC = 0.98). Performance are much poorer for the out-of-distribution weak-vs-non-binder classification (AUC = 0.69), as shown in Fig. 1b; note that for this task we switched labels, *i*.*e*. weak binders are presented as strong ones and non-binders are deemed as weak, otherwise we would get AUC = 0.31).

We display in Fig. 1c the 2d projections of the embedded sequences in the last layer of our architecture (before linear classification) using tSNE. The tSNE visualization shows that the predictive power of the model depends on the location of the out-of-distribution cluster in the feature space compared to the in-distribution data. The out-of-distribution sequence distribution has a large overlap with the one of weak binders, which makes discrimination hard. Conversely, in this feature space, strong binders are well separated from the rest and hence the decision boundary learned over strong-vs-weak is efficient even against out-of-distribution data.

###### Natural CDR3*β*

To assess if the results derived in the controlled framework of synthetic data also holds for natural TCRs, we consider the epitope `ELAGIGILTV`, which is 2 mutations away from the primary epitope `AAGIGILTV` expressed on the surface of Melanoma-cancer responsible cells (***Skipper et al., 1999)***. Our dataset includes 2,082 sequences of CDR3*β* experimentally labelled as binders to this epitope. We train the CNN model to distinguish peptide-specific sequences from bulk ones. We then select as out-of-distribution sequences (i) CDR3*β*s that positively bind the epitope `EAAGIGILTV`, one mutation away from the wildtype (WT) `ELAGIGILTV`; (ii) we also select CDR3*β* binding a very different peptide having Levenshtein distance 8 from our WT, namely VQELYSPIFLIV. We expect that our model be predictive for out-of-distribution specificity predictions for the first epitope and not for the second one. Results confirm this guess with values of AUC equal to, respectively, 0.79 and 0.54. Similarly to the case of synthetic data in Fig. 1c, this difference in performance is visualized in Fig. 2 by the distinct locations of the corresponding sequence distributions in the feature space of the last embedding layer of our classifier.

**Appendix 4—figure 2.**
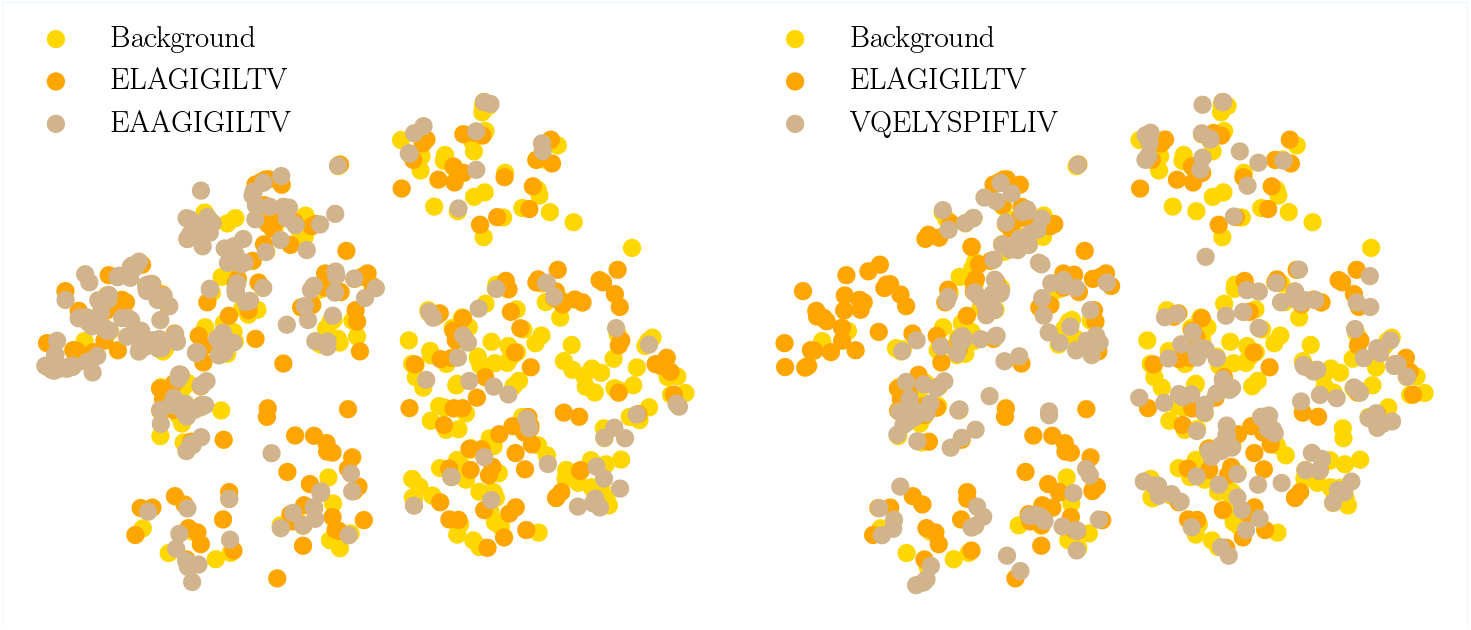
Out-of-distribution performances are related to the distribution of features. tSNE visualization of the feature vectors (in the second-last layer of the classifier architecture) of out-of-distribution data for a model trained over the WT epitope `ELAGIGILTV`, with a balanced dataset containing 1500 CDR3*β* per class. Top: As the WT epitope, `EAAGIGILTV` is responsible for the Melanoma cancer and thus is targeted by TCRs sharing similar features. Bottom: the VQELYSPIFLIV peptide is 8 mutations away from the WT and is involved in SARS-CoV-2 infections. Our model predictions are reliable in the first epitope (AUC = 0.79), and are not for the second peptide (AUC = 0.54). The tSNE plots support the claim that out-of-distribution specificity predictions drop when the feature vectors of the test data are far away from the training ones.

#### Performances of pan-specific models strongly depend on the choice of the out-of-distribution data

More generally, as we already discussed, the performance degradation is related to how much we move away from the in-distribution data. We report the table of out-of-distribution performances as a function of in-to out-epitopes Levenshtein distance (see Tab. 1).

**Appendix 4—table 1.** Pan-specific prediction scores for out-of-distribution tasks. The AUC (second column) and ACC (fourth column) scores are averaged across many out-of-distribution epitopes, with balance restored through pan-specific generation. Out-of-distribution CDR3*β* are grouped according to the Levenshtein distance of their associated epitope to the closest one in the training dataset (first column). Predictions worsen as peptides get further away from the ones in the training dataset. The third and fifth columns show the standard deviations of the scores.

| Dist. | AUC | $\sigma(\text{AUC})$ | ACC | $\sigma(\text{ACC})$ |
| --- | --- | --- | --- | --- |
| 1-3 | 0.68 | 0.05 | 0.65 | 0.05 |
| 4-6 | 0.63 | 0.10 | 0.61 | 0.06 |
| 7-9 | 0.62 | 0.09 | 0.60 | 0.07 |
| 10-11 | 0.49 | 0.20 | 0.52 | 0.11 |

## Appendix 5

### Hyperparameter tuning in pan-specific models

In Fig. 4 of the main text we show results for a specific value of the threshold size 𝒢, above which peptide-specific classes are not augmented. The choice of the threshold value can affect the performances and it is task-dependent. Here, we report in Fig. 1 results of AUC scores before and after data augmentation through the pan-specific model has been applied. Undersampling the populated classes is done down to size 5000 CDR3*β* sequences. As we can see, despite quantitative values depend on 𝒢, the picture confirms the overall concept that generative methods of peptide-specific CDR3*β* sequences help specificity predictions, particularly for under represented classes (small triangles).

**Appendix 5—figure 1.**
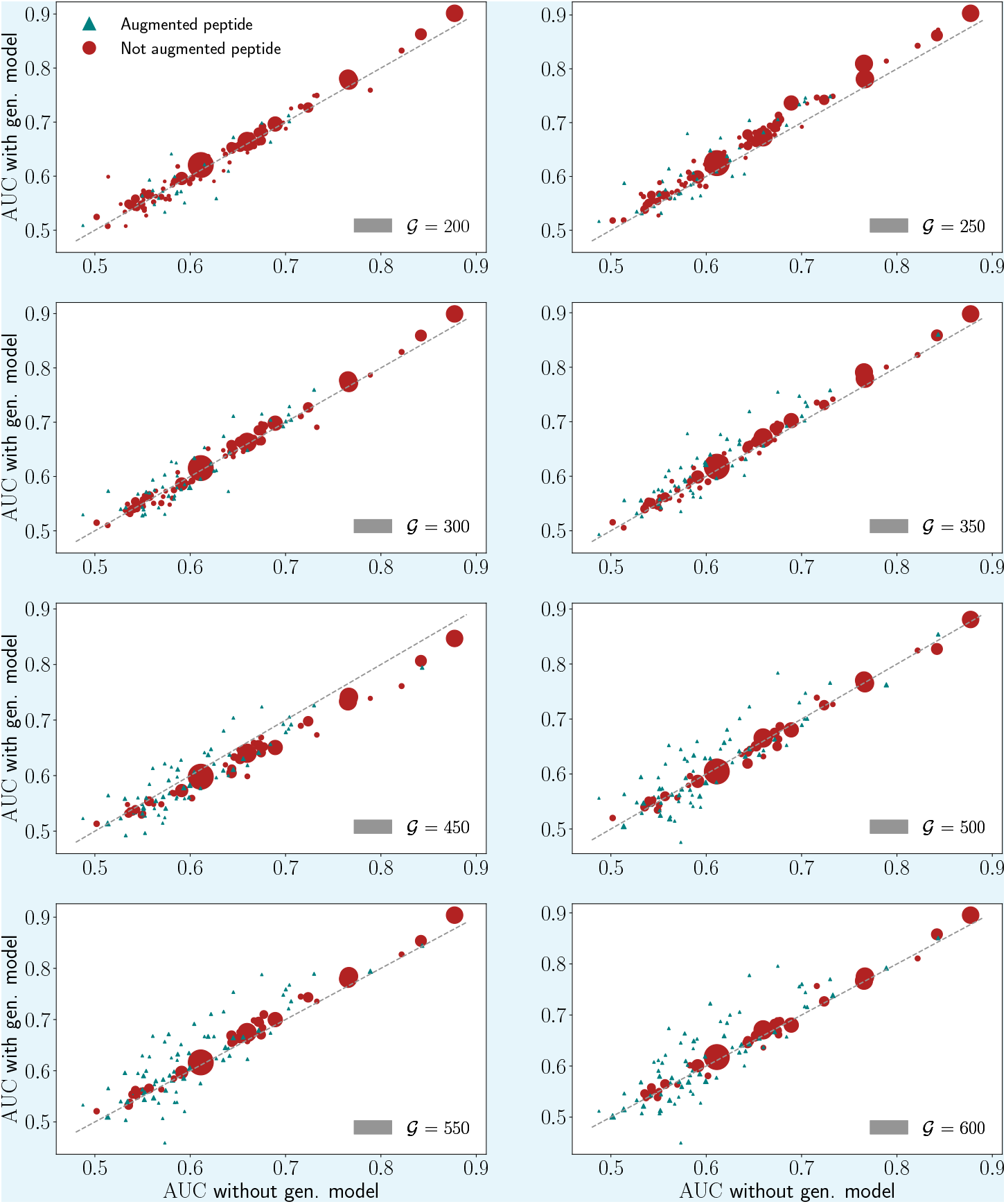
AUC scores dependencies on the hyperparameter choice *M*_*S*_. We report results of numerical experiments for the TCR-peptide binding prediction task, comparing AUC performances before and after the generative model has been used (x and y axis, respectively). The data point size is proportional to the size of natural peptide-specific sequences in the dataset. All the training parameters are fixed for all experiments; we rescale the number of training epochs with the training dataset size so that for each experiment we minimize the loss function exactly the same number of times: this factors out all elements, but the dependence of performances on the threshold value 𝒢.

## Appendix 6

### Details on Synthetic Lattice Protein dimer data

In the text we considered Lattice Proteins (LPs) as a fully controlled setting, where the ground truth distribution of data is known and its properties are tunable *ad hoc*. LPs consist of a computationally tractable model introduced to study the folding and binding properties of proteins (***Li et al., 1996; Mirny and Shakhnovich, 2001)***. A LP monomer is defined as a self-avoiding path over a 3 × 3 × 3 lattice cube, whose conformation defines a structure *S*. There are 𝒩 = 103, 406 distinct structures (up to global symmetries) on the cube. The probability that a sequence **v** of *L* = 27 amino acids folds into structure *S* is

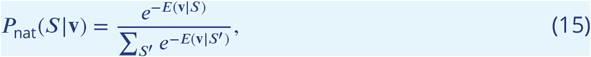

where the sum runs over a representative subset of all distinct structures. The energy of the sequence in a structure, *E*(**v**|*S*), is given by

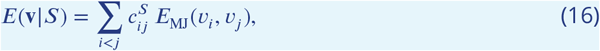

where *c*^*S*^ is the contact map of the structure (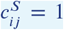if *i, j* are in contact, 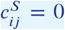 otherwise) and *E*_MJ_ is the Miyazawa-Jernigan energy matrix, a proxy for the effective interaction energy between pairs of amino acids.

Sequence data for LP dimers are obtained from ***Loffredo et al. (2023)***, where two monomer sequences **v**_1_, **v**_2_ – folded in, respectively, structures *S*_1_, *S*_2_ – form a dimeric complex via the interaction energy

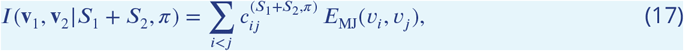

where the sum runs over all sites of both structures. The index *π* in (17) labels a specific orientation of the interaction. In analogy with (15), the probability that the sequences **v**_1_, **v**_2_ fold into the dimer *S*_1_ + *S*_2_ is

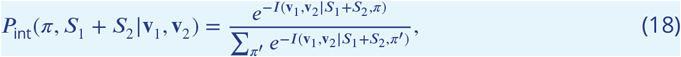

where the sum runs over all possible orientations. Given two structures *S*_1_, *S*_2_, sequences are collected through MCMC dynamics by accepting or rejecting a mutation at each evolution step based on the total probability

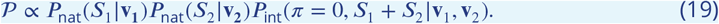

The MSAs are collected at steps *t* = 0, *t* = 100, *t* = 500 and represent non binder, weak and strong binder dimers used in this work. We refer the interested reader to ***Loffredo et al. (2023)*** for an extensive presentation of the LP model and details on how to obtain sequence data. In particular to generate the synthetic dataset used for the pan–specific model we select 12 pairs of structures and for each of them collect an MSA of binding and non binding sequences with the same length to ensure balance. The pairs of structures entering the dataset used in this work have the following ids: 1100 +1701; 249 +7801; 2789 +7511; 333 +794; 3412 + 9422; 456 + 259; 514 + 3894; 5809 + 6682; 9432 + 8754.

The out-of-distribution data for close structures is collected by first finding the closest structures to those used in the in-distribution dataset by minimizing the mean energy in (16)over the in-distribution binder MSA across all the structures available in ***Jacquin et al. (2016)***; ***Loffredo et al. (2023)***. We then generate the MSA for the selected close structure pair (2237 + 304), and this constitutes the close out-of-distribution dataset. Next we pick a random pair of structures and collect a MSA that will be our not close out-of-distribution dataset.

## Appendix 7

### Comparison of peptide-vs pan-specific generative models

Here we provide more details on the data augmentation in the pan-specific framework. In fact, it can be achieved in two ways.

1. By analogy with the peptide-specific case, we can design a single generative model that is trained on varied peptide and CDR3*β* pairs of sequences and produces new pairs. Intuitively, as the number of distinct CDR3*β*s dwarfs the one of epitopes, we expect a pan-specific generative model to first learn how to cluster groups of TCR sequences based on their epitope labels, then to learn the CDR3*β*s distribution within each group. However, such a model could suffer from group imbalance, and would learn effectively only the most-represented instances of epitopes and TCRs.
2. An alternative approach consists in training, separately, a generative model for each peptide-specific class of CDR3*β*s, so that the group imbalance present in the dataset is completely factored out. This second approach is however computationally demanding, as its running time increases linearly with the number of epitopes. In addition, overfitting could be an issue for peptide-specific groups with very little data.

As for the classifier, we closely follow the CNN architecture used in ***Montemurro et al. (2021)***, where two convolutional layers process separately the CDR3*β* and the peptide sequence: the resulting feature vectors are then concatenated to ultimately output the positive or negative binding prediction, see Model architecture for supervised predictions in the main text.

To assess the impact of group imbalance and the performance of the two approaches above, we first build an auxiliary dataset from the pan-specific dataset in ***Grazioli et al. (2022)*** by retaining only CDR3*β*s binding to five selected epitopes (`AMFWSVPTV`, `ELAGIGILTV`, `FLYNLLTRV`, `GLCTLVAML`, `LLWNGPMAV`), for a total of 9,000 datapoints. Fig. 1 reports the AUC and ACC scores obtained over each peptide and over the aggregate dataset comprising all five peptide-specific groups when CDR3*β* sequence data have been balanced with one global pan-specific generative model or with multiple peptide-specific ones. In the latter case, we have enlarged the CDR3*β* sequence space of the two most under represented group epitopes – `AMFWSVPTV` and `ELAGIGILTV`. We observe that predictive performances are comparable between the two approaches, suggesting that the pan-specific generative approach is not heavily impacted by group imbalance, and is capable of adequately clustering and modeling each peptide-associated group of TCR sequences.

This robustness stems from our sampling protocol, in which the generative model is carefully initialized with training sequences (Fig. 2). Upon sampling, the landscape defined by the inferred probability distribution is explored in the proximity of peptide-specific region associated to the initial sequence. This procedure prevents the generative model from jumping towards other peptide-specific groups, which can have much stronger overall weight due to group imbalance. Gibbs sampling schemes that start from randomly chosen sequence pairs preferably falls within such groups, *e*.*g*. peptides with less than 250 binders in the training dataset are on average generated four times less frequently than peptides with more.

**Appendix 7—figure 1.**
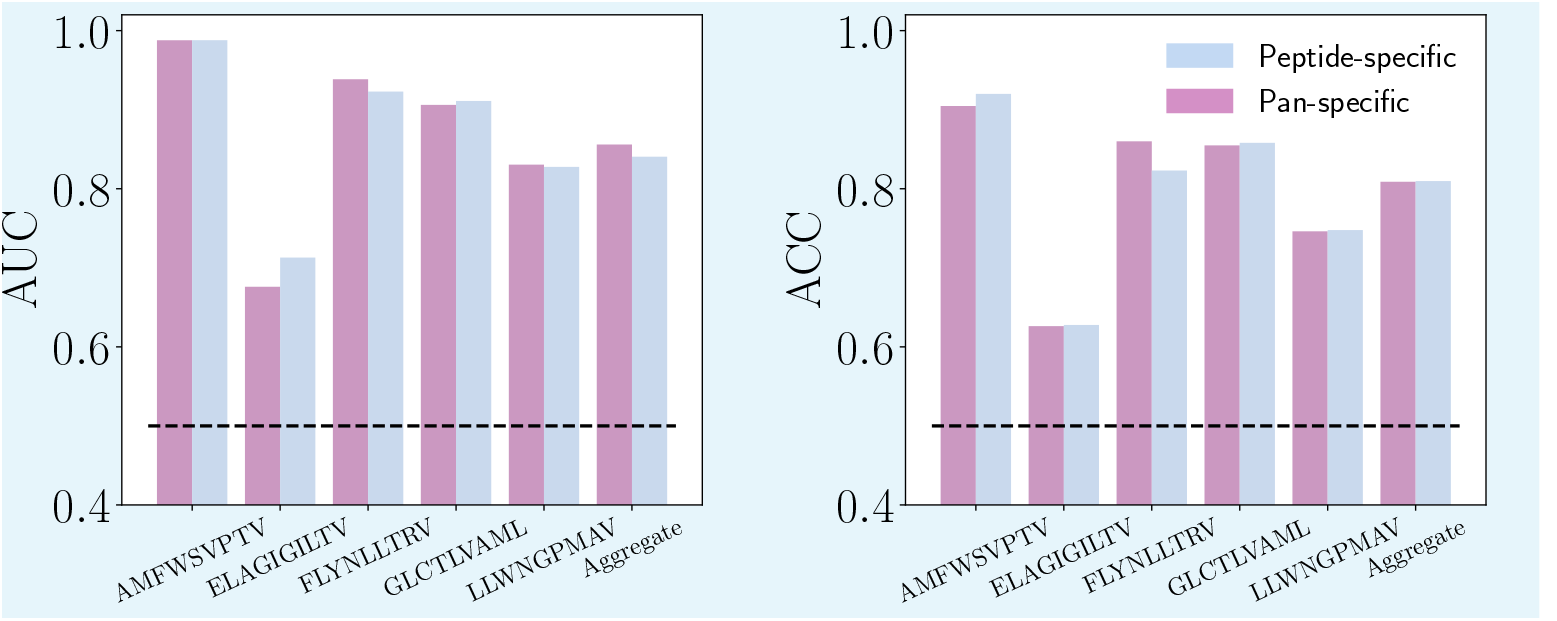
Performance change based on peptide-vs pan-specific generative model. Here we.report AUC and ACC scores for a subset of epitopes in the cases where each peptide class in augmented using a peptide-specific generative model or a global one. Given our sampling protocols, the two approaches are equivalent in terms of performance scores while the pan-specific remains computationally advantageous over the peptide-specific approach.

## Appendix 8

### Unsupervised classification

The pipeline explained in Results relies on the generative power of an unsupervised ML model trained on the positive examples and used to augment and balance the dataset, successively passed to a supervised classifier for training. To better understand how performances depend on the unsupervised and supervised steps, we report below predictions made by the unsupervised model alone. For simplicity, we consider hereafter the case of binary classification. By fixing a threshold on the scores, we can discriminate between positive and negative examples. This simple procedure is expected to be sub-optimal, as the unsupervised model is trained on the positive class alone and has no knowledge about the distribution of negative examples, which is used only to fix the threshold maximizing the accuracy on the two classes. Thus, a decision boundary based on the score of this model does not necessarily aligns with the separating surface of the two classes (see App. 2 for more details on the geometry of the classification problem).

In Fig. 1, we report the histograms of the scores assigned by the unsupervised model trained on positive examples of different peptide-CDR3*β* pairs, randomly chosen among the ones in Fig. 4 of the main text. The threshold (gray dashed line) is fixed by maximizing the accuracy on a validation set composed of positive and negative examples of the given peptide. Performances are consistently lower than the ones obtained using both the unsupervised model (to augment the data) and the CNN (to classify).

**Appendix 8—figure 1.**
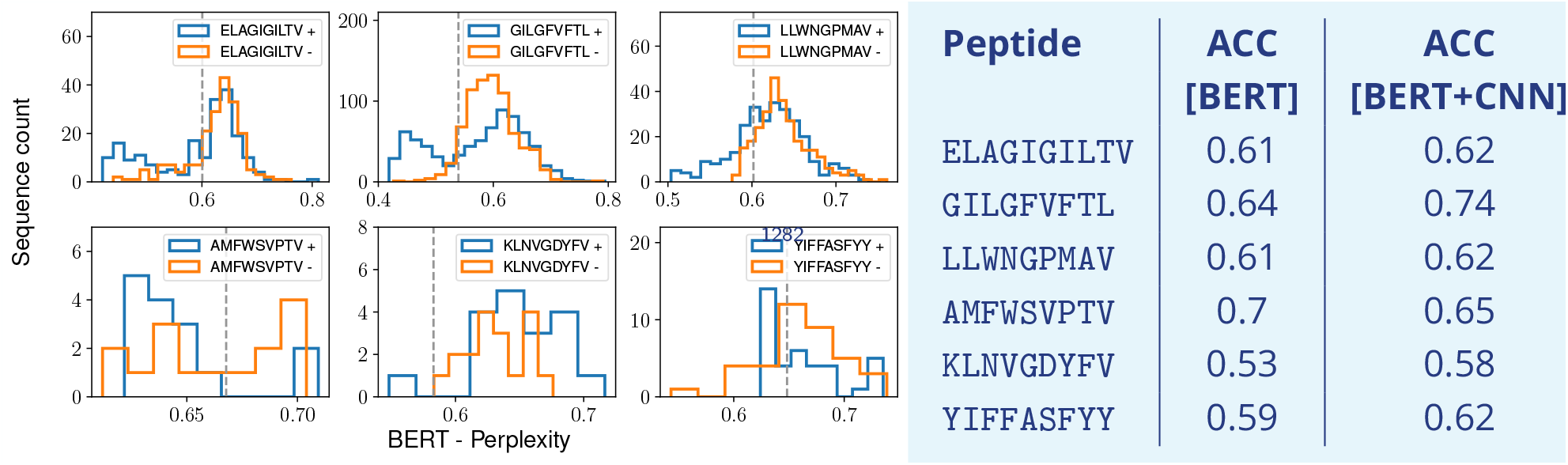
Unsupervised vs. supervised classification. **(Left)** Histograms of the scores assigned by the unsupervised model to balanced test sets of in-distribution epitopes; lower perplexity corresponds to better score. The gray dashed line locates the threshold obtained by maximizing the accuracy of prediction on positive and negative data. Epitopes in the top row are frequent ones and do not get enlarged during training of the supervised classifier. **(Right)** Comparison of the accuracy scores between unsupervised classification only ([BERT]) and the full pipeline ([BERT+CNN]) in Fig. 2 for the same peptides as in the left panel. Values in the column [BERT+CNN] can be obtained from Fig. 4 of the main text.

## References

Aguirre-López F, Franz S, Pastore M. Random features and polynomial rules. SciPost Phys. 2025; 18:039. https://scipost.org/10.21468/SciPostPhys.18.1.039, doi: 10.21468/SciPostPhys.18.1.039.

Ai Q, Wang P, He L, Wen L, Pan L, Xu Z. Generative Oversampling for Imbalanced Data via Majority-Guided VAE. In: Ruiz F, Dy J, van de Meent JW, editors. Proceedings of The 26th International Conference on Artificial Intelligence and Statistics, vol. 206 of Proceedings of Machine Learning Research PMLR; 2023. p. 3315–3330. https://proceedings.mlr.press/v206/ai23a.html.

Akbar R, Robert PA, Weber CR, Widrich M, Frank R, Pavlovic M, Scheffer L, Chernigovskaya M, Snapkov I, Slabodkin A, Mehta BB, Miho E, Lund-Johansen F, Andersen JT, Hochreiter S, Haff IH, Klambauer G, Sandve GK, and VG. In silico proof of principle of machine learning-based antibody design at unconstrained scale. mAbs. 2022; 14(1):2031482. http://dx.doi.org/10.1080/19420862.2022.2031482, doi: 10.1080/19420862.2022.2031482, pMID: 35377271.

Ansari M, White AD. Learning peptide properties with positive examples only. Digital Discovery. 2024; 3:977–986. http://dx.doi.org/10.1039/D3DD00218G, doi: 10.1039/D3DD00218G.

Bagaev DV, Vroomans RMA, Samir J, Stervbo U, Rius C, Dolton G, Greenshields-Watson A, Attaf M, Egorov ES, Zvyagin IV, Babel N, Cole DK, Godkin AJ, Sewell AK, Kesmir C, Chudakov DM, Luciani F, Shugay M. VDJdb in 2019: database extension, new analysis infrastructure and a T-cell receptor motif compendium. Nucleic Acids Research. 2019 10; 48(D1):D1057–D1062. http://dx.doi.org/10.1093/nar/gkz874, doi: 10.1093/nar/gkz874.

Bietti A, Mairal J. On the Inductive Bias of Neural Tangent Kernels. In: Wallach H, Larochelle H, Beygelzimer A, d’Alché-Buc F, Fox E, Garnett R, editors. Advances in Neural Information Processing Systems, vol. 32 Curran Associates, Inc.; 2019. https://proceedings.neurips.cc/paper/2019/file/c4ef9c39b300931b69a36fb3dbb8d60e-Paper.pdf.

Bravi B, Balachandran VP, Greenbaum BD, Walczak AM, Mora T, Monasson R, Cocco S. Probing T-cell response by sequence-based probabilistic modeling. PLOS Computational Biology. 2021 09; 17(9):1–27. https://doi.org/10.1371/journal.pcbi.1009297, doi: 10.1371/journal.pcbi.1009297.

Bravi B, Di Gioacchino A, Fernandez-de Cossio-Diaz J, Walczak AM, Mora T, Cocco S, Monasson R. A transferlearning approach to predict antigen immunogenicity and T-cell receptor specificity. eLife. 2023 sep; 12:e85126. http://dx.doi.org/10.7554/eLife.85126, doi: 10.7554/eLife.85126.

Chaudhuri K, Ahuja K, Arjovsky M, Lopez-Paz D. Why does Throwing Away Data Improve Worst-Group Error? In: Krause A, Brunskill E, Cho K, Engelhardt B, Sabato S, Scarlett J, editors. Proceedings of the 40th International Conference on Machine Learning, vol. 202 of Proceedings of Machine Learning Research PMLR; 2023. p. 4144–4188. https://proceedings.mlr.press/v202/chaudhuri23a.html.

Chen C, Kyathanahally SP, Reyes M, Merkli S, Merz E, Francazi E, Hoege M, Pomati F, Baity-Jesi Producing plankton classifiers that are robust to dataset shift. Limnology and Oceanography: Methods. 2025; 23(1):39–66. https://aslopubs.onlinelibrary.wiley.com/doi/abs/10.1002/lom3.10659, doi: 10.1002/lom3.10659.

Cheng Z, Zhou S, Guan J. Computationally predicting protein-RNA interactions using only positive and unlabeled examples. Journal of Bioinformatics and Computational Biology. 2015; 13(03):1541005. http://dx.doi.org/10.1142/S021972001541005X, doi: 10.1142/S021972001541005X.

Chizat L, Oyallon E, Bach F. On Lazy Training in Differentiable Programming. In: Wallach H, Larochelle H, Beygelzimer A, d’Alché-Buc F, Fox E, Garnett R, editors. Advances in Neural Information Processing Systems, vol. 32 Curran Associates, Inc.; 2019. https://proceedings.neurips.cc/paper/2019/file/ae614c557843b1df326cb29c57225459-Paper.pdf.

Croce G, Bobisse S, Moreno DL, Schmidt J, Guillame P, Harari A, Gfeller D. Deep learning predictions of TCRepitope interactions reveal epitope-specific chains in dual alpha T cells. Nature Communications. 2024 Apr; 15(1):3211. http://dx.doi.org/10.1038/s41467-024-47461-8, doi: 10.1038/s41467-024-47461-8.

Dandi Y, Stephan L, Krzakala F, Loureiro B, Zdeborová L. Universality laws for Gaussian mixtures in generalized linear models. In: Oh A, Naumann T, Globerson A, Saenko K, Hardt M, Levine S, editors. Advances in Neural Information Processing Systems, vol. 36 Curran Associates, Inc.; 2023. p. 54754–54768.

Dash P, Fiore-Gartland AJ, Hertz T, Wang GC, Sharma S, Souquette A, Crawford JC, Clemens EB, Nguyen THO, Kedzierska K, La Gruta NL, Bradley P, Thomas PG. Quantifiable predictive features define epitope-specific T cell receptor repertoires. Nature. 2017 Jul; 547(7661):89–93. http://dx.doi.org/10.1038/nature22383, doi: 10.1038/nature22383.

Dean J, Emerson RO, Vignali M, Sherwood AM, Rieder MJ, Carlson CS, Robins HS. Annotation of pseudogenic gene segments by massively parallel sequencing of rearranged lymphocyte receptor loci. Genome Medicine. 2015 Nov; 7(1):123. http://dx.doi.org/10.1186/s13073-015-0238-z, doi: 10.1186/s13073-015-0238-z.

Del Giudice, P, Franz, S, Virasoro, M A. Perceptron beyond the limit of capacity. J Phys France. 1989; 50(2):121–134. http://dx.doi.org/10.1051/jphys:01989005002012100, doi: 10.1051/jphys:01989005002012100.

Deng L, Ly C, Abdollahi S, Zhao Y, Prinz I, Bonn S. Performance comparison of TCR-pMHC prediction tools reveals a strong data dependency. Frontiers in Immunology. 2023; 14. https://www.frontiersin.org/journals/immunology/articles/10.3389/fimmu.2023.1128326, doi: 10.3389/fimmu.2023.1128326.

Dens C, Laukens K, Bittremieux W, Meysman P. The pitfalls of negative data bias for the T-cell epitope specificity challenge. Nature Machine Intelligence. 2023 Oct; 5(10):1060–1062. http://dx.doi.org/10.1038/s42256-023-00727-0, doi: 10.1038/s42256-023-00727-0.

Devlin J, Chang MW, Lee K, Toutanova K, BERT: Pre-training of Deep Bidirectional Transformers for Language Understanding; 2019. https://arxiv.org/abs/1810.04805.

Dietrich R, Opper M, Sompolinsky H. Statistical Mechanics of Support Vector Networks. Phys Rev Lett. 1999 04; 82:2975–2978. https://link.aps.org/doi/10.1103/PhysRevLett.82.2975, doi: 10.1103/PhysRevLett.82.2975.

Douzas G, Bacao F. Effective data generation for imbalanced learning using conditional generative adversarial networks. Expert Systems with Applications. 2018; 91:464–471. https://www.sciencedirect.com/science/article/pii/S0957417417306346, doi: 10.1016/j.eswa.2017.09.030.

Emerson RO, DeWitt WS, Vignali M, Gravley J, Hu JK, Osborne EJ, Desmarais C, Klinger M, Carlson CS, Hansen JA, Rieder M, Robins HS. Immunosequencing identifies signatures of cytomegalovirus exposure history and HLA-mediated effects on the T cell repertoire. Nature Genetics. 2017 May; 49(5):659–665. http://dx.doi.org/10.1038/ng.3822, doi: 10.1038/ng.3822.

Fernández A, García S, Galar M, Prati RC, Krawczyk B, Herrera F. Learning from Imbalanced Data Sets. Springer Cham; 2018. doi: 10.1007/978-3-319-98074-4.

Forman G, Scholz M. Apples-to-apples in cross-validation studies: pitfalls in classifier performance measurement. SIGKDD Explor Newsl. 2010 Nov; 12(1):49–57. http://dx.doi.org/10.1145/1882471.1882479, doi: 10.1145/1882471.1882479.

Fotouhi S, Asadi S, Kattan MW. A comprehensive data level analysis for cancer diagnosis on imbalanced data. Journal of Biomedical Informatics. 2019; 90:103089. https://www.sciencedirect.com/science/article/pii/S1532046418302302, doi: 10.1016/j.jbi.2018.12.003.

de G Matthews AG, Hron J, Rowland M, Turner RE, Ghahramani Z. Gaussian Process Behaviour in Wide Deep Neural Networks. In: International Conference on Learning Representations; 2018. https://openreview.net/forum?id=H1-nGgWC-.

Gerace F, Loureiro B, Krzakala F, Mézard M, Zdeborová L. Generalisation error in learning with random features and the hidden manifold model. Journal of Statistical Mechanics: Theory and Experiment. 2021 12; 2021(12):124013. http://dx.doi.org/10.1088/1742-5468/ac3ae6, doi: 10.1088/1742-5468/ac3ae6.

Ghoreyshi ZS, George JT. Quantitative approaches for decoding the specificity of the human T cell repertoire. Frontiers in Immunology. 2023; Volume 14 - 2023. https://www.frontiersin.org/journals/immunology/articles/10.3389/fimmu.2023.1228873, doi: 10.3389/fimmu.2023.1228873.

Gielis S, Moris P, Bittremieux W, De Neuter N, Ogunjimi B, Laukens K, Meysman P. Detection of Enriched T Cell Epitope Specificity in Full T Cell Receptor Sequence Repertoires. Frontiers in Immunology. 2019; Volume 10 - 2019. https://www.frontiersin.org/journals/immunology/articles/10.3389/fimmu.2019.02820, doi: 10.3389/fimmu.2019.02820.

Grazioli F, Mösch A, Machart P, Li K, Alqassem I, O’Donnell TJ, Min MR. On TCR binding predictors failing to generalize to unseen peptides. Frontiers in Immunology. 2022; 13. https://www.frontiersin.org/journals/immunology/articles/10.3389/fimmu.2022.1014256, doi: 10.3389/fimmu.2022.1014256.

Haque MM, Skinner MK, Holder LB. Imbalanced Class Learning in Epigenetics. Journal of Computational Biology. 2014; 21(7):492–507. http://dx.doi.org/10.1089/cmb.2014.0008, doi: 10.1089/cmb.2014.0008, pMID: 24798423.

Henikoff S, Henikoff JG. Amino acid substitution matrices from protein blocks. Proceedings of the National Academy of Sciences. 1992; 89(22):10915–10919. https://www.pnas.org/doi/abs/10.1073/pnas.89.22.10915, doi: 10.1073/pnas.89.22.10915.

Idrissi BY, Arjovsky M, Pezeshki M, Lopez-Paz D. Simple data balancing achieves competitive worst-group-accuracy. In: Schölkopf B, Uhler C, Zhang K, editors. Proceedings of the First Conference on Causal Learning and Reasoning, vol. 177 of Proceedings of Machine Learning Research PMLR; 2022. p. 336–351. https://proceedings.mlr.press/v177/idrissi22a.html.

Isacchini G, Walczak AM, Mora T, Nourmohammad A. Deep generative selection models of T and B cell receptor repertoires with soNNia. Proceedings of the National Academy of Sciences. 2021; 118(14):e2023141118. https://www.pnas.org/doi/abs/10.1073/pnas.2023141118, doi: 10.1073/pnas.2023141118.

Jacot A, Gabriel F, Hongler C. Neural Tangent Kernel: Convergence and Generalization in Neural Networks. In: Bengio S, Wallach H, Larochelle H, Grauman K, Cesa-Bianchi N, Garnett R, editors. Advances in Neural Information Processing Systems, vol. 31 Curran Associates, Inc.; 2018. https://proceedings.neurips.cc/paper/2018/file/5a4be1fa34e62bb8a6ec6b91d2462f5a-Paper.pdf.

Jacquin H, Gilson A, Shakhnovich E, Cocco S, Monasson R. Benchmarking Inverse Statistical Approaches for Protein Structure and Design with Exactly Solvable Models. PLOS Computational Biology. 2016 05; 12(5):1–18. http://dx.doi.org/10.1371/journal.pcbi.1004889, doi: 10.1371/journal.pcbi.1004889.

Jensen MF, Nielsen M. NetTCR 2.2 - Improved TCR specificity predictions by combining pan- and peptide-specific training strategies, loss-scaling and integration of sequence similarity. ELife. 2023 Oct; 12:RP93934. http://dx.doi.org/10.1101/2023.10.12.562001, doi: 10.1101/2023.10.12.562001.

Klinger M, Pepin F, Wilkins J, Asbury T, Wittkop T, Zheng J, Moorhead M, Faham M. Multiplex Identification of Antigen-Specific T Cell Receptors Using a Combination of Immune Assays and Immune Receptor Sequenc-ing. PLOS ONE. 2015 10; 10(10):1–21. http://dx.doi.org/10.1371/journal.pone.0141561, doi: 10.1371/jour-nal.pone.0141561.

Krawczyk B, Galar M, Lukasz Jelen, Herrera F. Evolutionary undersampling boosting for imbalanced classifi-cation of breast cancer malignancy. Applied Soft Computing. 2016; 38:714–726. https://www.sciencedirect.com/science/article/pii/S1568494615005815, doi: 10.1016/j.asoc.2015.08.060.

Kyathanahally SP, Hardeman T, Merz E, Bulas T, Reyes M, Isles P, Pomati F, Baity-Jesi M. Deep Learning Clas-sification of Lake Zooplankton. Frontiers in Microbiology. 2021; 12. https://www.frontiersin.org/journals/microbiology/articles/10.3389/fmicb.2021.746297, doi: 10.3389/fmicb.2021.746297.

Lee J, Sohl-dickstein J, Pennington J, Novak R, Schoenholz S, Bahri Y. Deep Neural Networks as Gaussian Pro-cesses. In: International Conference on Learning Representations; 2018. https://openreview.net/forum?id=B1EA-M-0Z.

Li H, Helling R, Tang C, Wingreen N. Emergence of Preferred Structures in a Simple Model of Protein Folding. Science. 1996; 273(5275):666–669. https://www.science.org/doi/abs/10.1126/science.273.5275.666, doi: 10.1126/science.273.5275.666.

Li M, Wu Z, Wang W, Lu K, Zhang J, Zhou Y, Chen Z, Li D, Zheng S, Chen P, Wang B. Protein-Protein Interaction Sites Prediction Based on an Under-Sampling Strategy and Random Forest Algorithm. IEEE/ACM Transactions on Computational Biology and Bioinformatics. 2022; 19(6):3646–3654. doi: 10.1109/TCBB.2021.3123269.

Lobo JM, Jiménez-Valverde A, Real R. AUC: a misleading measure of the performance of predictive distribution models. Global Ecology and Biogeography. 2008; 17(2):145–151. https://onlinelibrary.wiley.com/doi/abs/10.1111/j.1466-8238.2007.00358.x, doi: 10.1111/j.1466-8238.2007.00358.x.

Loffredo E, Vesconi E, Razban R, Peleg O, Shakhnovich E, Cocco S, Monasson R. Evolutionary dynamics of a lattice dimer: a toy model for stability vs. affinity trade-offs in proteins. Journal of Physics A: Mathematical and Theoretical. 2023 oct; 56(45):455002. https://dx.doi.org/10.1088/1751-8121/acfddc, doi: 10.1088/1751-8121/acfddc.

Loffredo E, Pastore M, Cocco S, Monasson R. Restoring balance: principled under/oversampling of data for optimal classification. In: Salakhutdinov R, Kolter Z, Heller K, Weller A, Oliver N, Scarlett J, Berkenkamp F, edi-tors. Proceedings of the 41st International Conference on Machine Learning, vol. 235 of Proceedings of Machine Learning Research PMLR; 2024. p. 32643–32670. https://proceedings.mlr.press/v235/loffredo24a.html.

Loureiro B, Gerbelot C, Cui H, Goldt S, Krzakala F, Mezard M, Zdeborová L. Learning curves of generic features maps for realistic datasets with a teacher-student model. In: Ranzato M, Beygelzimer A, Dauphin Y, Liang PS, Vaughan JW, editors. Advances in Neural Information Processing Systems, vol. 34 Cur-ran Associates, Inc.; 2021. p. 18137–18151. https://proceedings.neurips.cc/paper_files/paper/2021/file/9704a4fc48ae88598dcbdcdf57f3fdef-Paper.pdf.

Loureiro B, Sicuro G, Gerbelot C, Pacco A, Krzakala F, Zdeborová L. Learning Gaussian Mixtures with Generalized Linear Models: Precise Asymptotics in High-dimensions. In: Ranzato M, Beygelzimer A, Dauphin Y, Liang PS, Vaughan JW, editors. Advances in Neural Information Processing Systems, vol. 34 Cur-ran Associates, Inc.; 2021. p. 10144–10157. https://proceedings.neurips.cc/paper_files/paper/2021/file/543e83748234f7cbab21aa0ade66565f-Paper.pdf.

Lu T, Zhang Z, Zhu J, Wang Y, Jiang P, Xiao X, Bernatchez C, Heymach JV, Gibbons DL, Wang J, Xu L, Reuben A, Wang T. Deep learning-based prediction of the T cell receptor–antigen binding specificity. Nature Machine Intelligence. 2021 Oct; 3(10):864–875. http://dx.doi.org/10.1038/s42256-021-00383-2, doi: 10.1038/s42256-021-00383-2.

Mannelli SS, Gerace F, Rostamzadeh N, Saglietti L, Bias-inducing geometries: an exactly solvable data model with fairness implications; 2023. https://arxiv.org/abs/2205.15935.

Marouf M, Machart P, Bansal V, Kilian C, Magruder DS, Krebs CF, Bonn S. Realistic in silico generation and augmentation of single-cell RNA-seq data using generative adversarial networks. Nature Communications. 2020 Jan; 11(1):166. http://dx.doi.org/10.1038/s41467-019-14018-z, doi: 10.1038/s41467-019-14018-z.

Mason DM, Reddy ST. Predicting adaptive immune receptor specificities by machine learning is a data gen-eration problem. Cell Systems. 2024; 15(12):1190–1197. https://www.sciencedirect.com/science/article/pii/S2405471224003442, doi: 10.1016/j.cels.2024.11.008.

Meynard-Piganeau B, Feinauer C, Weigt M, Walczak AM, Mora T. TULIP: A transformer-based unsupervised language model for interacting peptides and T cell receptors that generalizes to unseen epitopes. Proceed-ings of the National Academy of Sciences. 2024; 121(24):e2316401121. https://www.pnas.org/doi/abs/10.1073/pnas.2316401121, doi: 10.1073/pnas.2316401121.

Meysman P, Barton J, Bravi B, Cohen-Lavi L, Karnaukhov V, Lilleskov E, Montemurro A, Nielsen M, Mora T, Pereira P, Postovskaya A, Martínez MR, de Cossio-Diaz JF, Vujkovic A, Walczak AM, Weber A, Yin R, Eug-ster A, Sharma V. Benchmarking solutions to the T-cell receptor epitope prediction problem: IMMREP22 workshop report. ImmunoInformatics. 2023; 9:100024. https://www.sciencedirect.com/science/article/pii/S2667119023000046, doi: 10.1016/j.immuno.2023.100024.

Ming Z, Chen X, Wang S, Liu H, Yuan Z, Wu M, Xia H. HostNet: improved sequence representation in deep neural networks for virus-host prediction. BMC Bioinformatics. 2023 Dec; 24(1):455. http://dx.doi.org/10.1186/s12859-023-05582-9, doi: 10.1186/s12859-023-05582-9.

Mirny L, Shakhnovich E. Protein Folding Theory: From Lattice to All-Atom Models. Annual Review of Biophysics. 2001; 30:361–396. https://www.annualreviews.org/content/journals/10.1146/annurev.biophys.30.1.361, doi: 10.1146/annurev.biophys.30.1.361.

Mirza B, Haroon D, Khan B, Padhani A, Syed TQ. Deep Generative Models to Counter Class Imbalance: A Model-Metric Mapping With Proportion Calibration Methodology. IEEE Access. 2021; 9:55879–55897. doi: 10.1109/ACCESS.2021.3071389.

Mondal AK, Singhal L, Tiwary P, Singla P, Ap P. Minority Oversampling for Imbalanced Data via Class-Preserving Regularized Auto-Encoders. In: Ruiz F, Dy J, van de Meent JW, editors. Proceedings of The 26th International Conference on Artificial Intelligence and Statistics, vol. 206 of Proceedings of Machine Learning Research PMLR; 2023. p. 3440–3465. https://proceedings.mlr.press/v206/mondal23a.html.

Montemurro A, Schuster V, Povlsen HR, Bentzen AK, Jurtz V, Chronister WD, Crinklaw A, Hadrup SR, Winther O, Peters B, Jessen LE, Nielsen M. NetTCR-2.0 enables accurate prediction of TCR-peptide binding by using paired TCRα and β sequence data. Communications Biology. 2021 Sep; 4(1):1060. http://dx.doi.org/10.1038/s42003-021-02610-3, doi: 10.1038/s42003-021-02610-3.

Nagano Y, Pyo AGT, Milighetti M, Henderson J, Shawe-Taylor J, Chain B, Tiffeau-Mayer A. Contrastive learning of T cell receptor representations. Cell Systems. 2025 Jan; 16(1). http://dx.doi.org/10.1016/j.cels.2024.12.006, doi: 10.1016/j.cels.2024.12.006.

Neal RM. In: Priors for Infinite Networks New York, NY: Springer New York; 1996. p. 29–53. http://dx.doi.org/10.1007/978-1-4612-0745-0_2, doi: 10.1007/978-1-4612-0745-0_2.

Nolan S, Vignali M, Klinger M, Dines JN, Kaplan IM, Svejnoha E, Craft T, Boland K, Pesesky MW, Gittelman RM, Snyder TM, Gooley CJ, Semprini S, Cerchione C, Nicolini F, Mazza M, Delmonte OM, Dobbs K, Carreño-Tarragona G, Barrio S, et al. A large-scale database of T-cell receptor beta sequences and binding associations from natural and synthetic exposure to SARS-CoV-2. Frontiers in Immunology. 2025; Volume 16 - 2025. https://www.frontiersin.org/journals/immunology/articles/10.3389/fimmu.2025.1488851, doi: 10.3389/fimmu.2025.1488851.

Pesce L, Krzakala F, Loureiro B, Stephan L. Are Gaussian Data All You Need? The Extents and Limits of Uni-versality in High-Dimensional Generalized Linear Estimation. In: Krause A, Brunskill E, Cho K, Engelhardt B, Sabato S, Scarlett J, editors. Proceedings of the 40th International Conference on Machine Learning, vol. 202 of Proceedings of Machine Learning Research PMLR; 2023. p. 27680–27708. https://proceedings.mlr.press/v202/pesce23a.html.

Pezzicoli FS, Ros V, Landes FP, Baity-Jesi M, Class Imbalance in Anomaly Detection: Learning from an Exactly Solvable Model; 2025. https://arxiv.org/abs/2501.11638.

Pham MDN, Nguyen TN, Tran LS, Nguyen QTB, Nguyen TPH, Pham TMQ, Nguyen HN, Giang H, Phan MD, Nguyen V. epiTCR: a highly sensitive predictor for TCR–peptide binding. Bioinformatics. 2023 04; 39(5):btad284. http://dx.doi.org/10.1093/bioinformatics/btad284, doi: 10.1093/bioinformatics/btad284.

Rana P, Sowmya A, Meijering E, Song Y. Imbalanced classification for protein subcellular localization with multilabel oversampling. Bioinformatics. 2022 12; 39(1):btac841. http://dx.doi.org/10.1093/bioinformatics/btac841, doi: 10.1093/bioinformatics/btac841.

Richardson E, Trevizani R, Greenbaum JA, Carter H, Nielsen M, Peters B. The receiver operating characteristic curve accurately assesses imbalanced datasets. Patterns. 2024; 5(6):100994. https://www.sciencedirect.com/science/article/pii/S2666389924001090, doi: 10.1016/j.patter.2024.100994.

Robert PA, Akbar R, Frank R, Pavlovic M, Widrich M, Snapkov I, Slabodkin A, Chernigovskaya M, Scheffer L, Smorodina E, Rawat P, Mehta BB, Vu MH, Mathisen IF, Prósz A, Abram K, Olar A, Miho E, Haug DTT, Lund-Johansen F, et al. Unconstrained generation of synthetic antibody–antigen structures to guide machine learn-ing methodology for antibody specificity prediction. Nature Computational Science. 2022 Dec; 2(12):845–865. http://dx.doi.org/10.1038/s43588-022-00372-4, doi: 10.1038/s43588-022-00372-4.

Sagawa S, Koh PW, Hashimoto TB, Liang P. Distributionally Robust Neural Networks for Group Shifts: On the Importance of Regularization for Worst-Case Generalization. CoRR. 2019; abs/1911.08731. http://arxiv.org/abs/1911.08731.

Saito T, Rehmsmeier M. The Precision-Recall Plot Is More Informative than the ROC Plot When Evaluating Binary Classifiers on Imbalanced Datasets. PLOS ONE. 2015 03; 10(3):1–21. http://dx.doi.org/10.1371/journal.pone.0118432, doi: 10.1371/journal.pone.0118432.

Sethna Z, Isacchini G, Dupic T, Mora T, Walczak AM, Elhanati Y. Population variability in the generation and selection of T-cell repertoires. PLOS Computational Biology. 2020 12; 16(12):1–21. http://dx.doi.org/10.1371/journal.pcbi.1008394, doi: 10.1371/journal.pcbi.1008394.

Sgarbossa D, Lupo U, Bitbol AF. Generative power of a protein language model trained on multiple sequence alignments. eLife. 2023 feb; 12:e79854. http://dx.doi.org/10.7554/eLife.79854, doi: 10.7554/eLife.79854.

Shugay M, Bagaev DV, Zvyagin IV, Vroomans RM, Crawford JC, Dolton G, Komech EA, Sycheva AL, Koneva AE, Egorov ES, Eliseev AV, Van Dyk E, Dash P, Attaf M, Rius C, Ladell K, McLaren JE, Matthews KK, Clemens EB, Douek DC, et al. VDJdb: a curated database of T-cell receptor sequences with known antigen speci-ficity. Nucleic Acids Research. 2017 09; 46(D1):D419–D427. http://dx.doi.org/10.1093/nar/gkx760, doi: 10.1093/nar/gkx760.

Sidhom JW, Larman HB, Pardoll DM, Baras AS. DeepTCR is a deep learning framework for revealing sequence concepts within T-cell repertoires. Nature Communications. 2021 Mar; 12(1):1605. http://dx.doi.org/10.1038/s41467-021-21879-w, doi: 10.1038/s41467-021-21879-w.

Sim MJW. TCRs and AI: the future is now. Nature Reviews Immunology. 2024 Jan; 24(1):3–3. http://dx.doi.org/10.1038/s41577-023-00974-7, doi: 10.1038/s41577-023-00974-7.

Skipper JCA, Gulden PH, Hendrickson RC, Harthun N, Caldwell JA, Shabanowitz J, Engelhard VH, Hunt DF, Slingluff Jr CL. Mass-spectrometric evaluation of HLA-A*0201-associated peptides identifies dominant nat-urally processed forms of CTL epitopes from MART-1 and gp100. International Journal of Cancer. 1999; 82(5):669–677. https://onlinelibrary.wiley.com/doi/abs/10.1002/%28SICI%291097-0215%2819990827%2982%3A5%3C669%3A%3AAID-IJC9%3E3.0.CO%3B2-%23, doi: 10.1002/(SICI)1097-0215(19990827)82:5<669::AID-IJC9>3.0.CO;2-\#.

Song H, Bremer BJ, Hinds EC, Raskutti G, Romero PA. Inferring Protein Sequence-Function Relationships with Large-Scale Positive-Unlabeled Learning. Cell Systems. 2021; 12(1):92–101.e8. https://www.sciencedirect.com/science/article/pii/S2405471220304142, doi: 10.1016/j.cels.2020.10.007.

Springer I, Besser H, Tickotsky-Moskovitz N, Dvorkin S, Louzoun Y. Prediction of Specific TCR-Peptide Binding From Large Dictionaries of TCR-Peptide Pairs. Frontiers in Immunology. 2020; Volume 11 - 2020. https://www.frontiersin.org/journals/immunology/articles/10.3389/fimmu.2020.01803, doi: 10.3389/fimmu.2020.01803.

Springer I, Tickotsky N, Louzoun Y. Contribution of T Cell Receptor Alpha and Beta CDR3, MHC Typing, V and J Genes to Peptide Binding Prediction. Frontiers in Immunology. 2021; Volume 12 - 2021. https://www.frontiersin.org/journals/immunology/articles/10.3389/fimmu.2021.664514, doi: 10.3389/fimmu.2021.664514.

Tickotsky N, Sagiv T, Prilusky J, Shifrut E, Friedman N. McPAS-TCR: a manually curated catalogue of pathology-associated T cell receptor sequences. Bioinformatics. 2017 05; 33(18):2924–2929. http://dx.doi.org/10.1093/bioinformatics/btx286, doi: 10.1093/bioinformatics/btx286.

Tubiana J, Cocco S, Monasson R. Learning protein constitutive motifs from sequence data. eLife. 2019 mar; 8:e39397. http://dx.doi.org/10.7554/eLife.39397, doi: 10.7554/eLife.39397.

Ursu E, Minnegalieva A, Rawat P, Chernigovskaya M, Tacutu R, Sandve GK, Robert PA, Greiff V. Training data composition determines machine learning generalization and biological rule discovery. bioRxiv. 2024; https://www.biorxiv.org/content/early/2024/06/19/2024.06.17.599333, doi: 10.1101/2024.06.17.599333.

Vita R, Mahajan S, Overton JA, Dhanda SK, Martini S, Cantrell JR, Wheeler DK, Sette A, Peters B. The Immune Epitope Database (IEDB): 2018 update. Nucleic Acids Research. 2018 10; 47(D1):D339–D343. https://doi.org/10.1093/nar/gky1006, doi: 10.1093/nar/gky1006.

Wan Z, Zhang Y, He H. Variational autoencoder based synthetic data generation for imbalanced learning. In: 2017 IEEE Symposium Series on Computational Intelligence (SSCI); 2017. p. 1–7. doi: 10.1109/SSCI.2017.8285168.

Wang A, Cho K, BERT has a Mouth, and It Must Speak: BERT as a Markov Random Field Language Model; 2019. https://arxiv.org/abs/1902.04094.

Wang C, Ding C, Meraz RF, Holbrook SR. PSoL: a positive sample only learning algorithm for finding non-coding RNA genes. Bioinformatics. 2006 08; 22(21):2590–2596. http://dx.doi.org/10.1093/bioinformatics/btl441, doi: 10.1093/bioinformatics/btl441.

Weber A, Pélissier A, Martínez MR, T cell receptor binding prediction: A machine learning revolution; 2023. https://arxiv.org/abs/2312.16594.

Williams C. Computing with Infinite Networks. In: Mozer MC, Jordan M, Petsche T, editors. Advances in Neu-ral Information Processing Systems, vol. 9 MIT Press; 1996. https://proceedings.neurips.cc/paper/1996/file/ae5e3ce40e0404a45ecacaaf05e5f735-Paper.pdf.

Wu KE, Yost K, Daniel B, Belk J, Xia Y, Egawa T, Satpathy A, Chang H, Zou J. TCR-BERT: learning the grammar of T-cell receptors for flexible antigen-binding analyses. In: Knowles DA, Mostafavi S, editors. Proceedings of the 18th Machine Learning in Computational Biology meeting, vol. 240 of Proceedings of Machine Learning Research PMLR; 2024. p. 194–229. https://proceedings.mlr.press/v240/wu24b.html.

Yadav S, Vora DS, Sundar D, Dhanjal JK. TCR-ESM: Employing protein language embeddings to predict TCR-peptide-MHC binding. Computational and Structural Biotechnology Journal. 2024 Dec; 23:165–173. http://dx.doi.org/10.1016/j.csbj.2023.11.037, doi: 10.1016/j.csbj.2023.11.037.

Yang P, Li XL, Mei JP, Kwoh CK, Ng SK. Positive-unlabeled learning for disease gene identification. Bioinfor-matics. 2012 08; 28(20):2640–2647. http://dx.doi.org/10.1093/bioinformatics/bts504, doi: 10.1093/bioinformat-ics/bts504.

Zhang N, Bevan MJ. CD8+ T Cells: Foot Soldiers of the Immune System. Immunity. 2011 Aug; 35(2):161–168. http://dx.doi.org/10.1016/j.immuni.2011.07.010, doi: 10.1016/j.immuni.2011.07.010.

Zhang W, Wang L, Liu K, Wei X, Yang K, Du W, Wang S, Guo N, Ma C, Luo L, Wu J, Lin L, Yang F, Gao F, Wang X, Li T, Zhang R, Saksena NK, Yang H, Wang J, et al. PIRD: Pan Immune Repertoire Database. Bioinformatics. 2019 08; 36(3):897–903. http://dx.doi.org/10.1093/bioinformatics/btz614, doi: 10.1093/bioinformatics/btz614.

Zieba M, Tomczak JM, Gonczarek A. RBM-SMOTE: Restricted Boltzmann Machines for Synthetic Minority Over-sampling Technique. In: Nguyen NT, Trawinski B, Kosala R, editors. Intelligent Information and Database Sys-tems Cham: Springer International Publishing; 2015. p. 377–386. doi: 10.1007/978-3-319-15702-3_37.

